# A model of the interactions between the FtsQLB and the FtsWI peptidoglycan synthase complex in bacterial cell division

**DOI:** 10.1101/2022.10.30.514410

**Authors:** Samuel J. Craven, Samson G. F. Condon, Alessandro Senes

**Author notes:** These authors contributed equally (Samuel J. Craven, Samson G. F. Condon).

## Abstract

In *Escherichia coli*, an important step in the divisome assembly pathway is the recruitment of the essential cell wall synthase complex FtsWI to the division site through interactions with the regulatory FtsQLB complex. Here, we investigate a key aspect of this recruitment by characterizing the structural organization of the FtsL-FtsW interaction. Mutations in the cytoplasmic and transmembrane regions of the two proteins result in cell division defects and loss of FtsW localization to division sites. We use these *in vivo* results to help validate the predicted interfaces from an AlphaFold2 model for the entire FtsQLBWI complex. Given the consistency between the predicted FtsQLBWI model and our current understanding of the structure and function of the complex, we further remodeled it, seeking insight into the potential structural transitions that may lead to activation of the FtsWI complex and PG synthesis. The model suggests that FtsLB serves as a support for FtsI, placing its periplasmic domain in an extended and possibly active conformation but it is also compatible with a proposed compact and possibly inactive conformation. Additionally, we reconfigure the model into an Fts[QLBWI]_2_ diprotomeric state, which suggests that FtsLB may act as a central hub during assembly of the PG synthesis machinery. Finally, we propose a possible role for FtsQ in activation of this machinery, potentially by acting as a gatekeeper for the interaction between the FtsL AWI region and FtsI. We propose that this gatekeeping function depends on a hinge next to the FtsLB CCD region, which has implications for the mechanisms behind the FtsLB *off*/*on* transition that is central to cell division regulation.

## Introduction

The bacterial growth cycle culminates in the splitting of the mother cell to form two genetically identical daughter cells, a process referred to as cell division or cytokinesis. Though cell division may appear relatively simple at first glance, multiple events must be properly coordinated in order to avoid potentially catastrophic mishaps like chromosomal damage or cell lysis. These events include establishment of the proper site of division, segregation of the newly replicated chromosomes into separate daughter cells, and finally breakage and reconstruction of the cell wall at the division site into a septum that will form the new poles of the daughter cells. These various aspects of cell division are coordinated through the activities of a multiprotein complex, referred to as the divisome, which assembles into a ring-like structure around the circumference of the cell at the site of division (1,2).

Although over 30 proteins participate in divisome function in *Escherichia coli*, the most central component is the tubulin homolog FtsZ (3). As the first essential divisome protein to localize to the division site (4), FtsZ establishes where the septum will form by polymerizing into protofilaments (5,6) that provide a scaffold for recruitment of the rest of the divisome (1). FtsZ protofilaments also regulate divisome dynamics through their “treadmilling” activity (7,8), in which concomitant growth and shrinkage from opposing ends of a filament result in unidirectional movement of the entire filament without movement of the individual FtsZ monomers. The spatiotemporal dynamics of various cell wall synthesis enzymes have been shown to correlate with FtsZ treadmilling, including enzymes that are critical for the reconstruction of the cell wall (7–9).

The cell wall of Gram-negative bacteria like *E. coli* consists of a single layer of peptidoglycan (PG), a meshwork of polysaccharide chains crosslinked by short peptides. Reorganization of the PG layer into a septum requires the coordination of numerous essential and nonessential enzymes that break down PG at the division site and synthesize new material perpendicular to the long axis of the cell (10,11). For synthesis of new PG material, polysaccharide chains are first polymerized from lipid II precursors bearing a pentapeptide-linked disaccharide (composed of *N*-acetylmuramic acid and *N*-acetylglucosamine) via the action of glycosyltransferase (GTase) enzymes (12). These nascent PG polymers are then crosslinked via their pentapeptides and integrated with the existing PG layer by the activity of transpeptidase (TPase) enzymes.

The primary GTase activity during cell division is provided by the multipass membrane protein FtsW (13,14), whereas the essential division TPase activity is provided by the monofunctional class B penicillin-binding protein FtsI (also called PBP3 in *E. coli*) (15–17). The interaction between FtsW and FtsI is well recognized, forming together the cognate FtsWI GTase-TPase pair responsible for the essential PG synthesis at the septum (18,19). Structurally, FtsW is an integral membrane protein with approximately half of its 414 residues contributing to its ten transmembrane (TM) helices (20). FtsW is the division-specific member of the recently described SEDS (shape, elongation, division, and sporulation) family of proteins (13,14), and it is a close homolog of RodA, which is responsible for the same enzymatic activity during cell elongation (13,21,22). FtsI is a single-pass membrane protein consisting of three major domains: the N-terminal transmembrane helix, which mediates its association with FtsW (19); the C-terminal TPase domain, which is the catalytic module; and an intervening pedestal domain, which is critical for the regulation of the activity of the FtsWI complex (23,24). FtsW and FtsI work along with the class A penicillin-binding protein PBP1b, a bifunctional enzyme that can catalyze both TPase and GTase reactions (25,26); however, PBP1B is nonessential for cell division.

The localization of FtsWI to midcell and its regulation are dependent on the membrane protein complex formed by FtsQ, FtsL, and FtsB (FtsQLB). FtsL and FtsB are single-pass membrane proteins of similar topology consisting of an N-terminal transmembrane helix followed by a coiled-coil domain in the periplasm. They assemble into a heterotetrameric FtsLB complex consisting of two FtsL and two FtsB subunits arranged into an extended helical configuration (27–29). The C-terminal, periplasmic tail of FtsB binds with high affinity to FtsQ (30–32), and this interaction is needed for the localization of FtsLB to midcell (33). In turn, FtsQLB recruits FtsWI to midcell primarily through an interaction between FtsW and the N-terminal, cytoplasmic tail of FtsL (34).

Recent advances in our understanding of cell division regulation have indicated that the FtsQLB interaction with FtsWI plays an important role in triggering the final constriction that completes cell division (35,36), an event that is initiated by accumulation FtsN at the division site (37). As schematically illustrated in Fig. 1, FtsN recognizes and binds regions in the septal cell wall where the existing PG layer is undergoing degradation, and this accumulation signals that new PG material can be inserted (38,39). Either indirectly or through a direct interaction, FtsN induces FtsQLB to switch from an *off* to *on* state, which is then communicated to the FtsWI synthesis machinery to activate PG remodeling at the septum (35,36). Initially, inactive FtsWI follows treadmilling FtsZ filaments around the septum; however, upon activation the complex is released from FtsZ and begins moving propelled by the forces generated by processive septal PG synthesis (9).

**Fig. 1.**
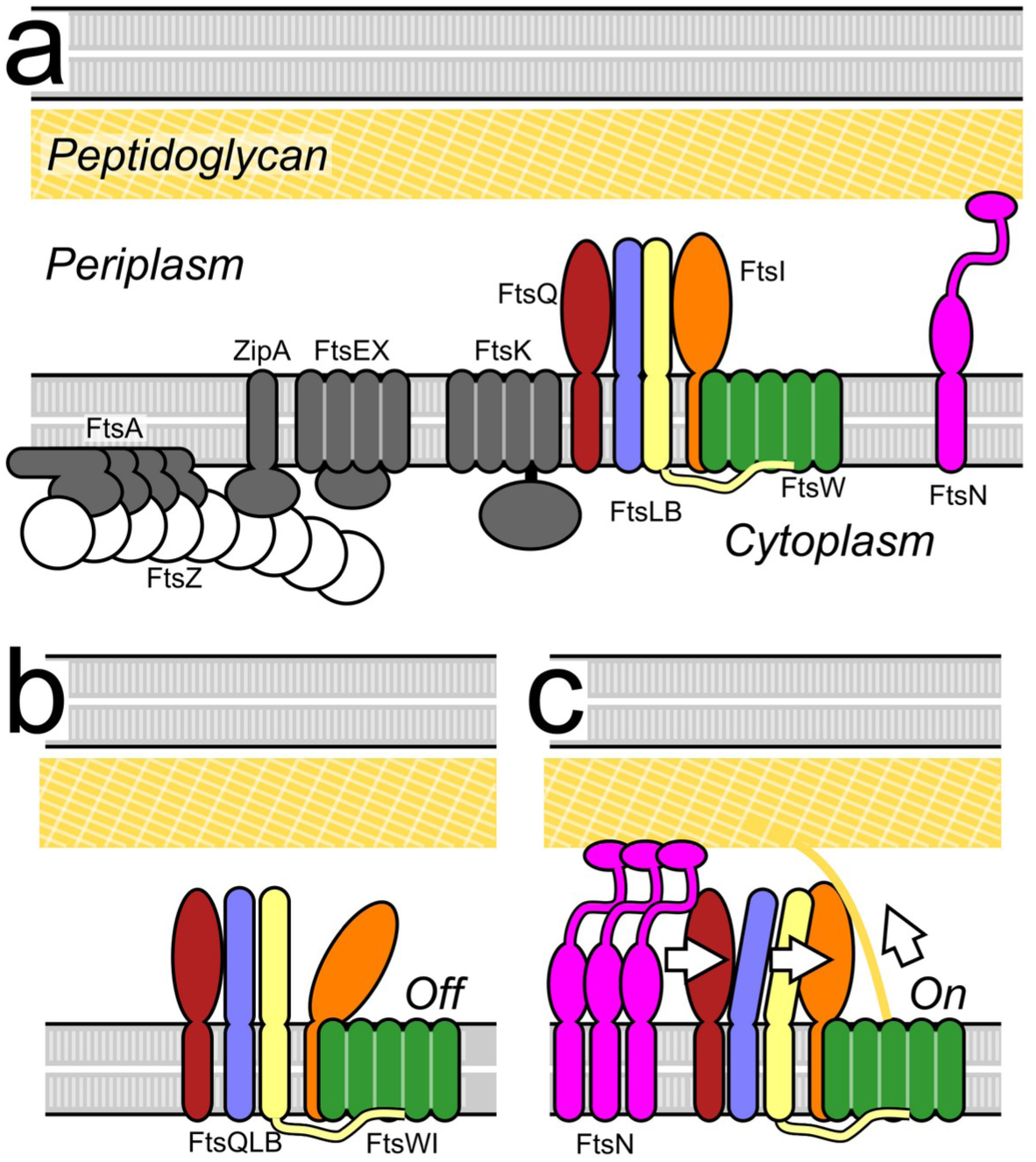
The current model for activation of cell division. a) Schematic representation of the essential components of the *E. coli* divisome. The polymeric ring formed by FtsZ establishes the site of division in coordination with other early components. The peptidoglycan synthase complex FtsWI is the main actor for the reconstruction of the cell wall, leading to the formation of a septum and, eventually, the poles of the nascent daughter cells. Its activation is tightly regulated by FtsN and FtsLB. b) In the current model, FtsWI is in complex with FtsQLB but is initially inactive. c) Accumulation of FtsN at midcell somehow triggers a conformational change in the FtsLB complex, which in turn triggers peptidoglycan synthesis likely by a direct interaction with FtsWI.

The hypothesis of FtsQLB being a regulator of FtsWI activity was proposed following the identification of gain-of-function mutations in a specific region of FtsL and FtsB that enable cell division to occur even in the absence FtsN (35,36). This region was named the Constriction Control Domain (CCD), and it consists of residues E_88_, N_89_, D_93_, and H_94_ in FtsL as well as A_55_, E_56_, and D_59_ in FtsB. Recently, a second critical region in FtsL was identified in which dominant-negative mutations can be made to prevent PG synthesis by FtsWI (40,41). This region has been termed the Activation of FtsWI domain (AWI; positions 82-84, 86-87, and 90), and it neighbors the CCD on the opposite helical face of FtsL without overlapping with it. The evidence suggest s that the *off/on* transition in FtsLB may involve a structural change around the CCD region that somehow enables the AWI region to activate FtsWI through a direct interaction with the periplasmic domain of FtsI (41) (Fig. 1c), although what constitutes the difference between the FtsQLB *off* and *on* states is currently unclear.

In order to obtain a better understanding of the structural and functional mechanisms behind activation of FtsWI by FtsQLB, we investigate here the interactions between these complexes. In particular, we focus on identifying the binding site between FtsL and FtsW, an interaction essential for FtsWI recruitment and, most likely, also critical for the stability of the FtsQLBWI complex and its proper functioning. We present an *in vivo* mutational study that identifies two sensitive regions of FtsL and FtsW (one in the juxtamembrane region on the cytoplasmic side of the proteins and the second involving the TM domains), in which mutation disrupts FtsW localization to the division site. We then analyze our results based on an AlphaFold2 (AF2) prediction of the FtsQLBWI complex. We find that the model produced by AF2 is consistent with our mutagenesis data and consensus observations from the literature and may thus hold important clues regarding the mechanism of FtsWI activation.

## Results and discussion

### A small region of the cytoplasmic tail proximal to the FtsL TM domain is critical for FtsW binding

In order to identify residues within the N-terminal, cytoplasmic tail of FtsL that are involved with the interaction with FtsW, we performed alanine-scanning mutagenesis *in vivo* using an FtsL depletion strain (MDG277) (33). By measuring the length distribution of cell populations expressing these FtsL variants, we assessed whether the mutants could properly support cell division or if they resulted in elongated cells due to cell division defects. To quantify the severity of cell division defects, we followed our previous method (28) in which we report the proportion of cells in the mutant population having lengths that exceed the 95^th^ percentile of the WT length distribution (dashed vertical line in the WT length distribution in Fig. 2b). The length distributions and representative cell images for all mutants tested are shown in supplementary Fig. S1 and Fig. S2. Protein expression levels for the various mutants were compared using western blot analysis to confirm that any defects seen were due to protein function instead of an inability to express to WT-like levels (supplementary Fig. S3a).

**Fig. 2.**
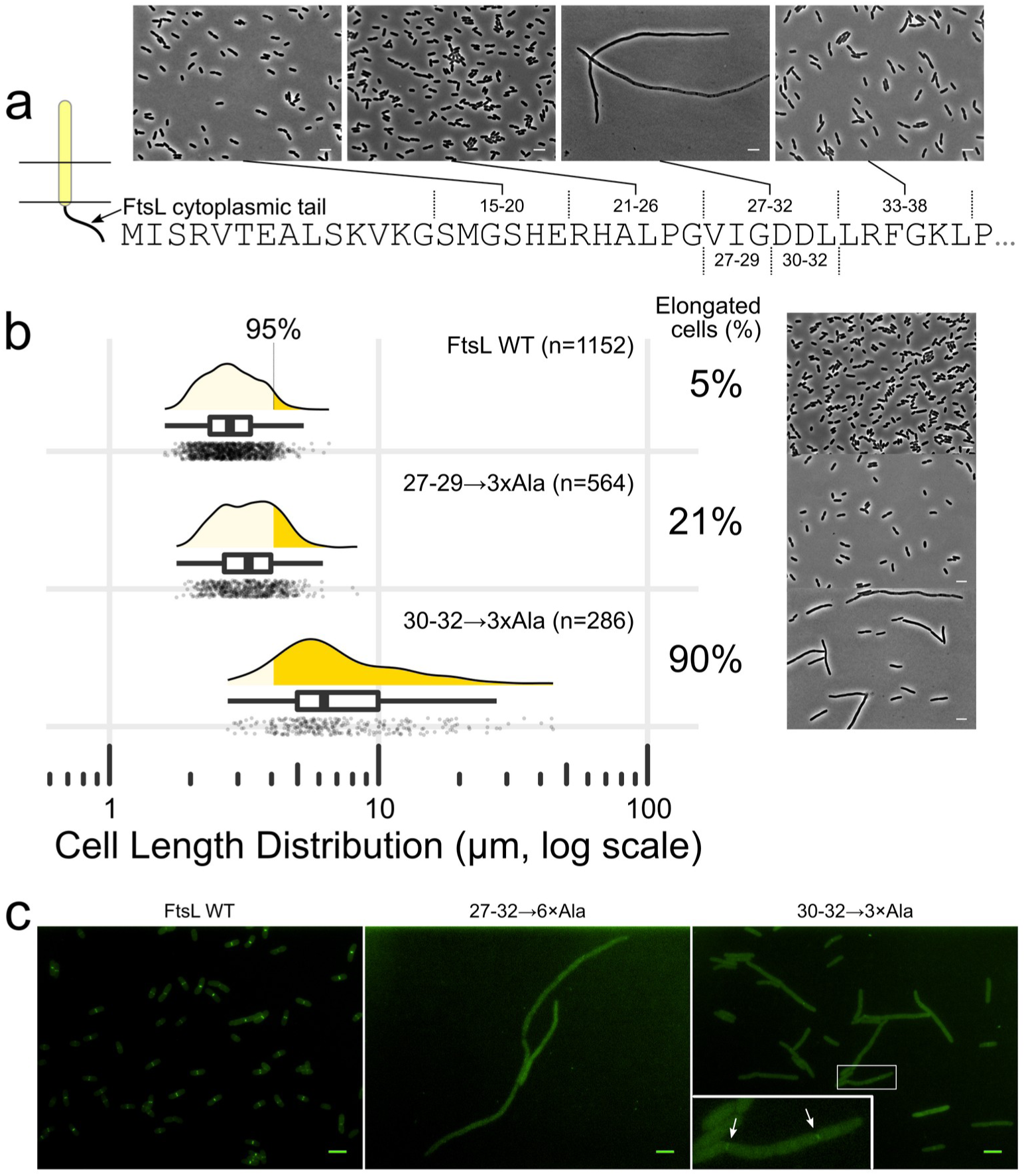
Mutations in the FtsL 27-32 region of the cytoplasmic tail disrupt FtsW localization. a) The essential region of the FtsL cytoplasmic tail was divided into four six-residue blocks, which were mutated to alanine (6×Ala) and tested for division and localization defects. Mutation of residues 27-32 produced a filamentous phenotype. b) When subdivided into two three-residue regions, the 27-29→3×Ala mutation produced a mild but notable elongation phenotype whereas the 30-32→3×Ala displayed 90% of cells that are longer than the 95^th^ percentile in the WT distribution (indicated as a dashed line in the WT histogram). c) The 27-32→6×Ala filaments do not display any notable GFP-FtsW localization foci. Foci are noticeably less frequent and less intense in 30-32→3×Ala (arrows in the magnified inset) than in cells expressing WT FtsL. Scale bars = 5 μm.

It was previously reported that truncation mutants of FtsL lacking the first 15 residues can complement depletion of the WT protein (34); therefore, the first third of the cytoplasmic tail is not required for FtsW binding and was excluded in our analysis. We instead focused on residues 15-38, which lead up to the approximate start of the transmembrane region of FtsL. We divided this region into four blocks of six residues each (residues 15-20, 21-26, 27-32, and 33-38; Fig. 2a), simultaneously mutating all residues in each block to alanine (6×Ala). The 6×Ala mutations in the 15-20, 21-26, and 33-38 blocks in the FtsL cytoplasmic tail resulted in relatively mild or moderate elongation defects; however, the 27-32→6×Ala mutation resulted in completely filamentous cells, identifying this segment as a candidate for FtsW binding (Fig. 2a).

We also introduced the 27-32→6×Ala mutants into another FtsL depletion strain (MDG254) expressing a GFP-labeled FtsW fusion protein (34) to test whether or not FtsW midcell localization was impacted by the FtsL mutants. As expected, we observed a loss of FtsW localization to division sites, as evidenced by a severe reduction in GFP foci formation (Fig. 2c). In these cells, the FtsL mutation resulted in a slightly less severe filamentation phenotype compared to MDG277, potentially due to overall increased FtsW levels resulting from GFP-FtsW being expressed in addition to native FtsW in MDG254. Nevertheless, this observation supports this region of FtsL being critical for FtsW binding.

To further narrow down which residues within the 27-32 region are important for division, we tested 3×Ala mutations in residues 27-29 and 30-32. As shown in Fig. 2b and supplementary Fig. S1, the 27-29→3×Ala mutation produced a mild division defect (21% elongated cells), whereas the 30-32→3×Ala mutation resulted in a much more severe defect (90%). When tested in strain MDG254, the FtsL 30-32→3×Ala mutation reproduced a similar elongation defect, but more importantly, GFP-FtsW foci were noticeably less frequent and less intense than in cells expressing WT FtsL (Fig. 2c). These data suggest that residues 30-32 are more critical for the interaction with FtsW. Additionally, the somewhat milder phenotype of the 30-32→3×Ala mutation compared to the 27-32→6×Ala mutation (severe elongation vs. complete filamentation, respectively) suggests that residues 27-29 likely also participate to some degree in the FtsL-FtsW binding site, which is supported by the mild elongation defect shown in Fig. 2b.

To further narrow down the role of each individual residue in the potential FtsL-FtsW binding region, single and double alanine mutations were introduced at positions 30-32. A mild division defect was observed for D31A (24% elongated cells), whereas D30A (12%) and L32A (14%) were even milder (supplementary Fig. S1). The D30A/L32A mutation resulted in a relatively mild defect (20%), whereas the D30A/D31A mutation and the D31A/L32A mutation resulted in moderate (39%) and severe (88%) defects, respectively. These data implicate D_31_ as a key residue in this potential FtsW-binding region of FtsL, though other residues (particularly L_32_) likely play a role in this interaction.

D_31_ may be involved in an important electrostatic interaction, such as a salt bridge, at the FtsL-FtsW interface, which would be lost in the D31A mutant. We hypothesized that a charge-reversal mutation would be repulsive against a corresponding positively charged region in FtsW, which would further destabilize the interaction interface. Indeed, a D31K mutation produced a severe elongation defect (90%), as opposed to the milder phenotype of D31A (24%) (supplementary Fig. S1), supporting the electrostatic interaction hypothesis. As a control, an analogous mutation introduced at position 30 (D30K) resulted in a very mild defect (10%), which further suggests that this position is not critically involved at the interaction interface. Overall, the data indicate that the region proximal to the TM domain of FtsL is the putative binding site for FtsW, with D _31_ and residues in its proximity being the most critical for this interaction.

### The putative FtsL-interaction site in FtsW is near the base of TM1 and TM10

The proposed FtsW-binding site in the FtsL cytoplasmic tail (residues 27-32) most likely interacts with residues exposed to the cytoplasmic face of FtsW, which consists of four loops connecting the transmembrane helices as well as the N- and C-terminal tails of FtsW, which are both cytoplasmic (Fig. 3). To identify the associated binding site in FtsW, we used a similar mutational strategy, under the hypothesis that mutations of residues critical for the interaction would likely result in defective recruitment of FtsW to the division site. We transformed plasmids containing variants of a functional GFP-FtsW fusion into the FtsW depletion strain EC912 (18), which contains a WT copy of FtsW under P_BAD_ control. This enabled us to simultaneously test for cell morphology and FtsW localization defects.

**Fig. 3.**
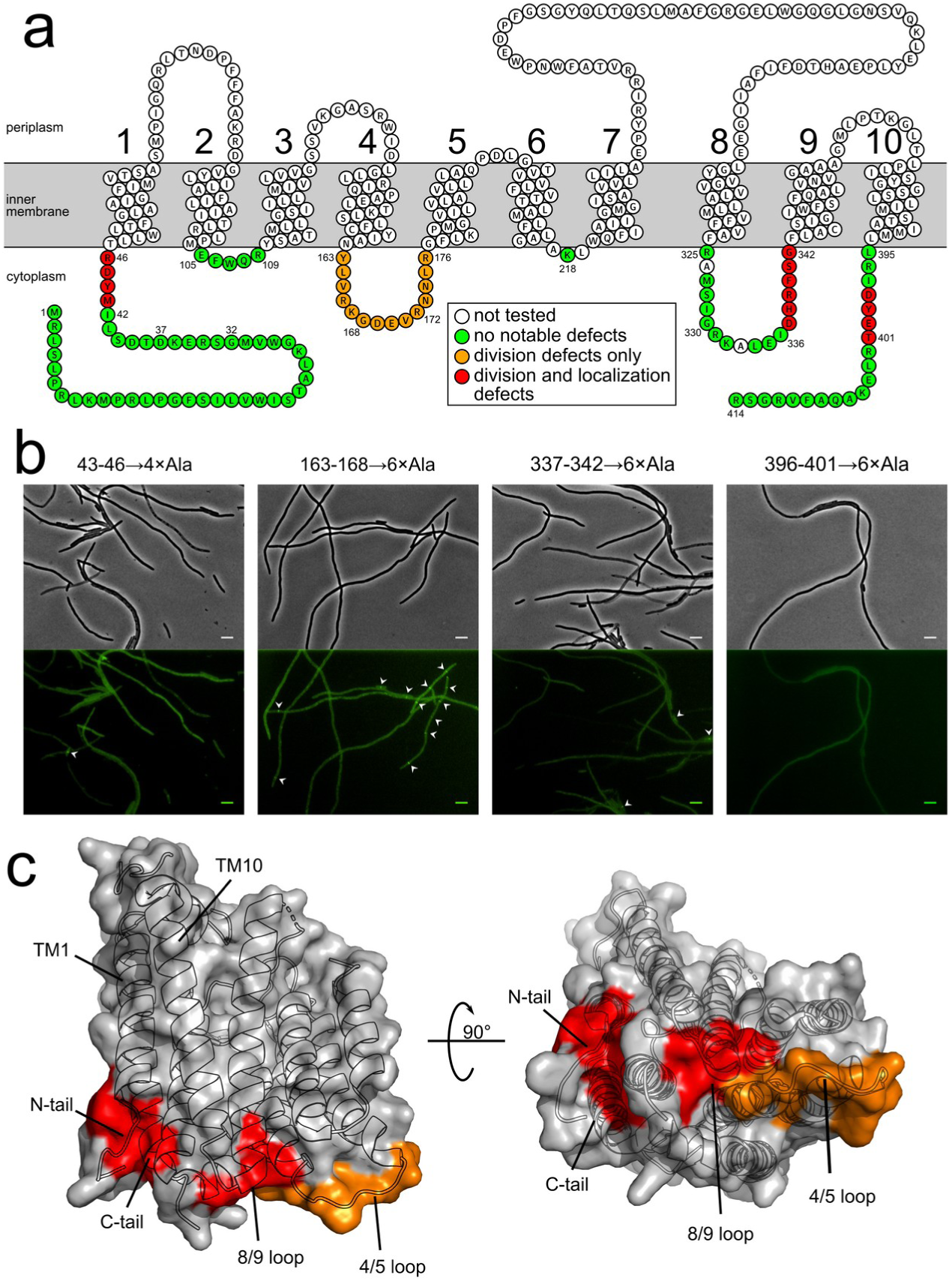
The mutations in FtsW that affect localization occur on distinct loops. a) Snake plot (Protter web application) representing the sequence and topology of *E. coli* FtsW (20). Small black numbers indicate relevant residue positions and large black numbers indicate the TM domains. The effects of different mutations (alanine-scanning or deletion) are color coded. Mutations that cause no or mild elongation defects are highlighted in green. Those that cause a stronger division defect but retain FtsW localization are coded in orange. Defective mutations that also resulted in reduced FtsW localization are coded in red. The latter are the likely candidates for the binding site of the FtsL tail. A list of the specific mutations tested with results are included in supplementary Table S1. Length distributions are included in supplementary Fig. S4. b) Representative cell images (phase-contrast on top; GFP fluorescence on bottom) for select FtsW mutations causing division defects. GFP foci are indicated by white arrows. Scale bars = 5 μm. c) Mapping of the sensitive mutations on the crystal structure of the FtsW SEDS family homolog RodA (PDB code 6BAR). The localization sensitive mutations (red) cluster on a distinct side of the cytoplasmic face of FtsW near the N- and C-terminal region of the protein.

Previous work indicated that FtsW residues 1-30 are not essential for function (42), but little was known about the importance of the remaining cytoplasm-facing residues. For this reason, we performed extensive mutagenesis on the entire cytoplasmic face of FtsW, by either truncation or mutation to alanine. The results are summarized graphically in Fig. 3 and in supplementary Table S1. All cell length distributions and representative cell images are provided in supplementary Fig. S4 and Fig. S5. GFP-FtsW expression levels for mutants lacking regular GFP foci in the cell images are compared using western blot analysis (supplementary Fig. S3a).

Complete truncations of either the N- or C-terminal FtsW tails (Δ1-46 or R396X, respectively, where “X” refers to a stop codon) resulted in severe elongation defects (88% and 91% elongation, respectively) as well as highly reduced GFP-FtsW localization at midcell. On the other hand, partial terminal truncations (Δ1-42 or R402X) resulted in mild elongation defects at most (7% and 13%, respectively) and displayed proper localization of GFP-FtsW at midcell. This suggested that loss of residues within the 43-46 or 396-401 regions caused the division defects seen in the full truncations of the tails, whereas the remainder of the tails are nonessential.

We then tested alanine substitutions in these regions (in full-length GFP-FtsW) to see if they could replicate the defects seen in the complete tail truncations. 4×Ala mutations in residues 43-46 and 398-401 resulted in the formation of severely elongated cells (92% and 79% elongation, respectively) lacking GFP-FtsW foci. Interestingly, separate 2×Ala substitutions in these regions resulted in cells with frequent GFP-FtsW foci and at most a moderate elongation phenotype (29% for 45-46→2×Ala), indicating that mutation of these smaller regions is not disruptive enough to cause severe division defects. Overall, these results indicate that large portions of the both the N-terminal and C-terminal tails of FtsW are disposable for FtsW localization or cell division in general, but the juxtamembrane residues on both N- and C-termini appear to be critical.

To identify whether the FtsL-binding site extends to any of the cytoplasmic loops of FtsW, we tested the phenotype of alanine mutations in each residue, with additional charge reversal mutations at select positions (either individually or in groups). As summarized in Fig. 3 (and shown in supplementary Fig. S4 and Table S1), many of these mutations resulted in no discernible defect in cell division or GFP-FtsW localization. A 6×Ala mutation in residues 337-342 of the 8/9 loop (the loop connecting transmembrane helices 8 and 9), however, resulted in a severe elongation phenotype (78%) with reduced GFP foci frequency (Fig. 3a,b). As indicated by Western blot analysis (supplementary Fig. S3b), this mutation resulted in decreased GFP-FtsW expression, suggesting that the elongation defect may be due to decreased levels of FtsW, though disruption of the FtsL-FtsW interaction is still a potential explanation. Subsequent 3×Ala mutations at positions 337-339 and 340-342 resulted in WT-like cells (5% and 2%, respectively) with regular GFP-FtsW foci, suggesting that mutation of multiple residues was needed to cause the defect resulting from the 337-342→6×Ala mutation.

Interestingly, multiple mutations throughout the 4/5 loop (residues 163-176) resulted in cell division defects (highlighted in orange in Fig. 3a), but in each case frequent GFP-FtsW foci could be seen throughout the elongated cells (Fig. 3b). These results suggest that the 4/5 loop variants retained the ability to bind FtsL and are more likely deficient in some other function of FtsW (for example, catalytic inactivation, inability to bind substrate, or an activation defect). One of these mutations (169-171→3×Ala) overlaps with a previously identified mutation (E170K) that was shown to disrupt cell division (43), though a clear mechanism was not proposed for the defect. More recently, however, this loop was proposed to play a role in the activation of FtsW activity through interaction with FtsA (44), which would be consistent with a cell division disruption phenotype without a defect in FtsW localization.

### The localization-sensitive mutations map on a distinct region of the cytoplasmic face of FtsW

We mapped the observed mutation-sensitive positions of FtsW over the crystal structure of *Thermus thermophilus* RodA (45), a close homolog of FtsW that performs a similar role in PG reconstruction during cell elongation (Fig. 3c). Because the terminal TM domains (TM1 and TM10) of RodA are in contact, the N- and C-terminal tails are also near each other in three-dimensional space. Additionally, these residues are also in close proximity to the 8/9 loop. Overall, the equivalent positions of the mutation-sensitive residues (FtsW positions 43-46, 337-342, and 396-401) form a continuous surface on the same half of the cytoplasmic face when mapped on RodA, as highlighted in red in Fig. 3c. This surface therefore represents a likely candidate for the FtsL-binding site on FtsW. In contrast, the mutation-sensitive region in the 4/5 loop (FtsW positions 163-176) is structurally contiguous to the 8/9 loop but occurs on the opposite end of the cytoplasmic face when mapped on RodA. Since this region impacts function but not localization of FtsW, this end of the cytoplasmic face is less likely to be part of the FtsL binding surface.

### An AlphaFold2 model of FtsQLBWI is consistent with the cytoplasmic binding site inferred from mutation

To rationalize the mutational data, we turned to protein structure prediction with AlphaFold2 (AF2) (46,47). Since FtsQLB and FtsWI are both assumed to form constitutive complexes (19,48), we modeled the FtsQLBWI complex instead of FtsL and FtsW alone. FtsLB assembles *in vitro* as a tetrameric (L_2_B_2_) complex (27,28), but FtsQLBWI was modeled in a single protomer configuration (i.e., with a single subunit for each protein) because AF2 does not predict a higher-oligomer configuration. However, the FtsLB dimer of AF2 superimposes well with half of our model of the complex in tetrameric configuration (29). The potential formation of a diprotomeric form of FtsQLBWI complex (Fts[QLBWI]_2_) is discussed in a later section.

The AF2 model of FtsQLBWI is illustrated in Fig. 4a. The individual chains align quite well with recently reported binary and ternary AF2 predictions (49). Despite the complex being over 1,000 residues in length and comprising five separate chains, the program generated five models that are very similar to one another with high confidence metrics (supplementary Fig. S6). The overall two-dimensional Predicted Alignment Error (PAE) plot of the complex indicates that in many regions there is high confidence in the positioning and orientation of the chains (supplementary Fig. S7a). Notably, the vast majority of PAEs for contacting residues between different chains in the FtsQLBWI complex are below 3 Å, suggesting confidence in the predicted protein:protein interfaces (supplementary Fig. S7b).

**Fig. 4.**
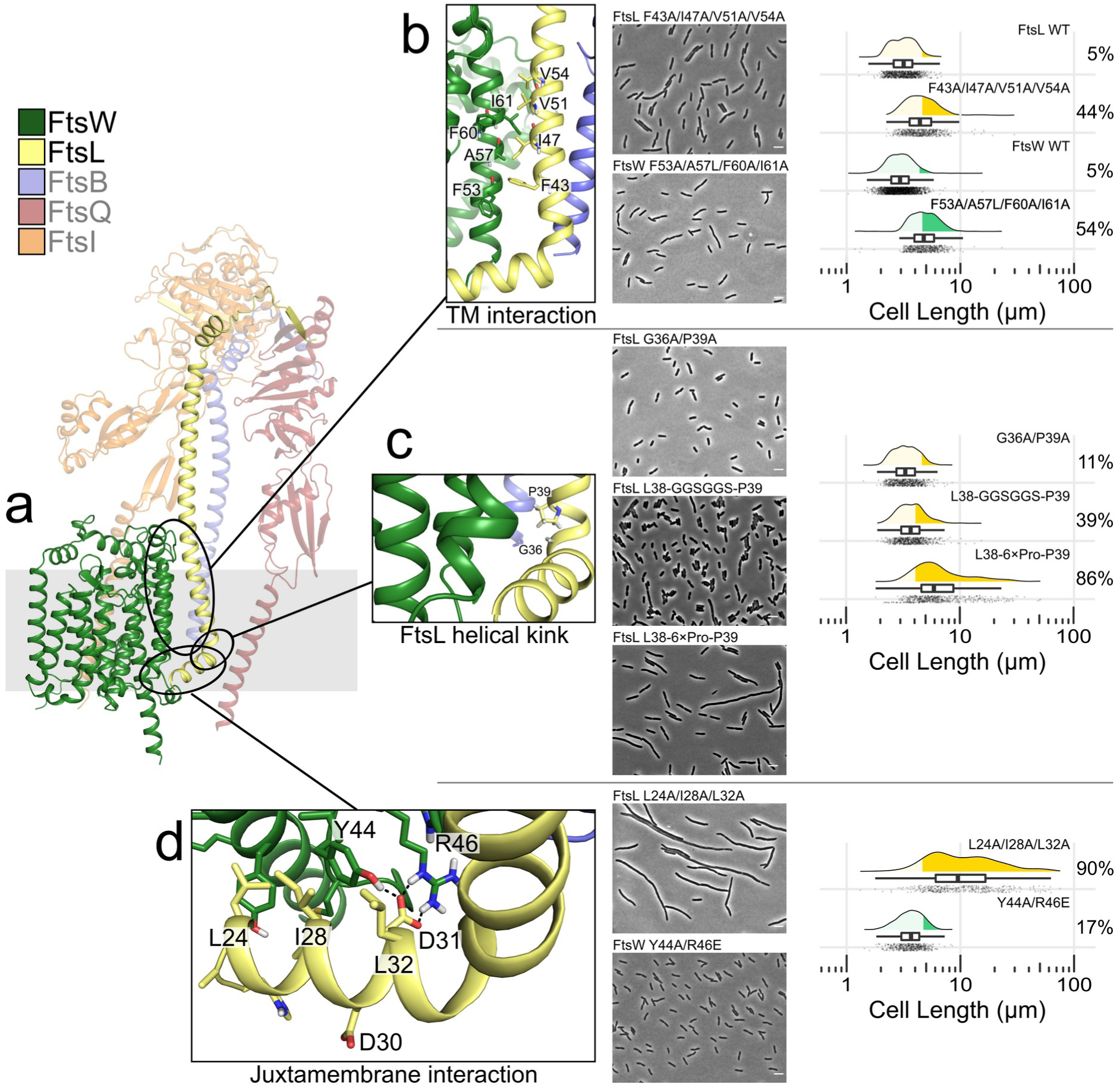
Mutation phenotypes interpreted over the AlphaFold2 structural prediction of the FtsQLBWI complex. a) View of the complex, with emphasis on the interaction between FtsL and FtsW. b) Details of the predicted interaction between FtsL TM and FtsW TM1. The four residues on each helix most involved in the interaction interface are marked. These were simultaneously mutated to Ala residues, producing significant elongation phenotypes, as illustrated. c) Mutation of the G36xxP39 motif at the base of FtsL TM results in a very mild elongation phenotype. Insertion of a floppy (GSSGSSG) or ridig (poly-Pro) sequence results in elongated cells. d) Details of the predicted interaction between the juxtamembrane helix of FtsL (residues 21-32) and the base of TM1 of FtsW. A predicted hydrogen bonding interaction between the critical FtsL residue D31 and Y44 and R46 of FtsW is highlighted. The non-critical D30 is fully exposed to solvent. The hydrophobic amino acids that contribute to packing of the juxtamembrane helix are marked. When these amino acids are substituted by Ala, a severe elongation phenotype is observed. Conversely, mutation of Y44 and R46 of FtsW produces only a mild phenotype. The model is available for download at the Dryad Digital Repository (DOI TBD).

The AF2 model is in overall good agreement with our experimental mutagenesis data. In the model, FtsL binds to FtsW through a short juxtamembrane α-helix corresponding to residues 24-35 (Fig. 4a,c), which is predicted with high confidence (PAE ~2-3 Å, supplementary Fig. S7c). This helix is parallel to the membrane plane, forming approximately a right angle with the FtsL TM helix. The juxtamembrane helix includes the region that we identified as the most sensitive to mutation (residues 27-32) along with part of the preceding region (21–26), which produced a mild phenotype (supplementary Fig. S1). Interestingly, the positions of the juxtamembrane helix that contact FtsW in the model (L_24_, I_28_, D_31_, and L_32_) include the two positions (D_31_ and L_32_) that displayed the most significant effects *in vivo*. Conversely, D_30_, whose mutation to Ala or Lys did not cause notable defects, is solvent exposed and not in contact with FtsW in the model.

Previous work identified a double mutation in two of the above-mentioned positions (L24K/I28K) that was proposed to weaken the FtsL-FtsW cytoplasmic interaction (41). The contact between these positions and FtsW in the AF2 model lends support to this hypothesis. Since the introduction of two charged residues at these positions could theoretically weaken the interaction indirectly, we decided to further validate the predicted cytoplasmic interactions by testing the phenotype of FtsL L24A/I28A/L32A, which represents the loss of the predicted hydrophobic interface of the FtsL juxtamembrane helix. This resulted in a dramatic elongation phenotype (90% elongated cells; Fig. 4a,c), which further supports the predicted interfacial location of those residues. We conclude that the experimental data relative to the FtsL juxtamembrane region strongly support the AF2 model.

The model is also in reasonable agreement with our mutagenesis data for FtsW, though the correspondence is not as direct as it is for FtsL. The most important finding is that the mutation-sensitive 43-46 region at the base of TM1 of FtsW (the 43-46→4×Ala mutation caused severe elongation and loss of FtsW localization) interacts directly with the key D_31_ residue of FtsL. Among the four FtsW residues in this region, R_46_ and Y_44_ are predicted to form hydrogen bonding interactions with FtsL D_31_ (Fig. 4c). In order to test the model, we mutated both FtsW positions in a variety of combinations (R46A or R46E plus Y44A or Y44F). Perhaps surprisingly, none of these combinations produced strong division or localization defects (supplementary Table S1), though Y44A/R46E (Fig. 4c) and Y44A/R46A did give mild elongation defects (17% and 13% elongated cells, respectively). Similarly, the aforementioned 45-46→2×Ala mutation produced a moderate phenotype (29%), suggesting that R46 is at least weakly involved with the FtsL interaction.

A region in the C-terminal tail of FtsW near the base of TM10 (positions 398-401) was also identified by our mutagenesis as a potential FtsL-binding site. This region is predicted by the models to be marginally involved at the protein-protein interface with the FtsL juxtamembrane α-helix. The only prominent contact identified by the models are between FtsW Y_398_ and the end of the FtsL juxtamembrane helix (Fig. 4). However, a double mutant 398-399→2×Ala displayed a WT-like phenotype in our analysis (supplementary Fig. S4), indicating that any interaction involving Y398 is not absolutely required for FtsL-FtsW binding. The final region shown by mutagenesis to affect FtsW localization, the 8/9 loop, was not highlighted by the AF2 model as a region for interaction with FtsL, which suggests that the severe elongation phenotype caused by the 396-401→6×Ala mutation could be solely due to decreased GFP-FtsW expression instead of reduced interaction with FtsL, as discussed previously.

The inconsistencies seen between the FtsW mutagenesis and modeling highlight the complexity of inferring protein-protein interaction from cellular phenotypes. Mutations of positions involved at interfaces do not necessarily produce sufficient destabilization to result in loss of localization. Conversely, it is also possible that mutation at positions that are not interfacial may have a distal effect on the interaction sites through structural rearrangements. Regardless of the noted inconsistencies, there is sufficient correspondence between the mutational data and the model to have confidence that the overall architecture of the FtsL-FtsW cytoplasmic interaction, with a primary binding site involving the juxtamembrane helix of FtsL and the 43-46 region at the base of TM1 of FtsW, is reasonably predicted.

### The interaction between the FtsL and FtsW TM domains contributes to stability of the complex

In addition to the cytoplasmic interaction between FtsL and FtsW, the AF2 model also predicts tight interactions within the TM and periplasmic regions of the proteins (Fig. 4a,d). As illustrated in supplementary Fig. S7c, the PAE contact map indicates a series of very high confidence (PAE ~1-2 Å) contacts between the TM helix of FtsL (positions 39-54) and both TM1 and the periplasmic 1/2 loop of FtsW (positions 50-78) as well as between the periplasmic juxtamembrane region of FtsL (positions 50-61) and positions 281-284 of the 7/8 loop of FtsW (commonly referred to as extracellular loop 4 or ECL4). The interaction between the TM domains occurs along a helical face of FtsL that is opposite from the previously established interface with FtsB (28,29), which results in a lack of interaction between FtsB and FtsW in the model. In addition, the FtsL-binding site on FtsW is distant from the well-established binding site for FtsI, which is recapitulated well by the model and formed by FtsW TM helices 8 and 9 (50).

To test the TM region binding site, we introduced mutations *in vivo* at this FtsL-FtsW interface. Since individual mutations are often not sufficient to produce significant effects, we simultaneously mutated all positions involved on the FtsL TM helix (F43A, I47A, V51A, and V54A) and, separately, all positions involved on FtsW-TM1 (F53A, A57L, F60A, and I61A). Both 4× mutant proteins resulted in moderate to severe elongation defects (44% elongated cells for the FtsL mutation and 54% for FtsW) (Fig. 4d). For both 4× mutations, GFP-FtsW foci were generally visible but only at sites of constriction, indicating that the changes caused defects in the localization of FtsW but, when FtsW could localize, the changes did not necessarily affect its function. The data support the notion that interaction in the membrane region between TM1 of FtsW and the TM domain of FtsL contributes to stabilizing the complex formation, further validating the AF2 model.

### The length and flexibility of an FtsL linker region may impact the interaction with FtsW

In the AF2 model, the juxtamembrane and transmembrane helices of FtsL are connected by a short, kinked linker segment centered on a G_36_xxP_39_ helix-breaker motif (Fig. 4c). As shown in supplementary Fig. S1, the 33-38→6×Ala mutation in the N-terminal, cytoplasmic tail of FtsL resulted in a moderate elongation defect (35% elongated cells), suggesting that this region has structural or functional relevance. One likely role is to facilitate the formation of the kink and enable proper relative orientation of the two FtsW-binding domains. Therefore, mutation of the G_36_xxP_39_ motif or deletions in this region would be expected to possibly cause cell division defects. We were also interested in probing whether additional flexibility in this region is tolerated in order to assess whether the cytoplasmic and transmembrane interaction regions form independent binding domains or if they are functionally or structurally coupled.

In the structural context of the model, P_39_ appears to have the dual purpose of inducing a helical break while also stabilizing the TM helix as a favorable N-capping residue, as observed, for example, in certain integrin molecules (51). We tested a P39N substitution, which lacks the helical break function but can act as an N-capping residue (52), as well as a P39A substitution, which would lead to the loss of both functions. Both mutations resulted in very mild cell division defects (11% elongated cells; supplementary Fig. S1). Similarly, mutation of G_36_ (G36A) alone or in combination with P39A also resulted WT-like (6% for G36A) or mild (11% for G36A/P39A, Fig. 4c) phenotypes, suggesting that the G_36_xxP_39_ motif is not essential for positioning the FtsW-binding site.

We expected deletion mutants in the juxtamembrane linker region of FtsL to be destabilizing based on the model conformation, since the ends of the juxtamembrane and/or transmembrane helices would have to unwind to connect the two segments. Accordingly, we found that deleting six residues (Δ33-38) resulted in a much more severe elongation phenotype compared to the corresponding 6×Ala mutation at that site (90% vs 35% elongated cells, respectively) (supplementary Fig. S1). Similarly, defect discrepancies were seen between deletion and Ala mutations in smaller regions, though the differences were less severe (20% and 26% for Δ33-35 and Δ36-38, respectively, compared to 13% and 22% for 33-35→3×Ala and 36-38→3×Ala, respectively). When combined with the mild phenotypes for the G_36_ and P_39_ mutations, these results indicate that the length of the linker region is more important than the specific residue identities.

We then tested whether the two FtsW-binding sites in FtsL need to be structurally coupled by increasing flexibility within the linker region, both by substitution (33-38→GGSGGS) as well as an insertion (L_38_-GGSGGS-P_39_) (Fig. 4c). Both mutations resulted in similar elongation defects (40% and 39% elongated cells, respectively), suggesting that the flexible sequence and not necessarily the added length caused the defect. Finally, to test the effect of rigidity, we introduced a poly-Pro region by substitution (33-38→6×Pro) and insertion (L_38_-6×Pro-P_39_). Poly-Pro adopts a typical helical structure that is extended and rigid in nature, and thus introduction of such a sequence would require a major rearrangement of the model’s structure. As expected, these drastic mutations resulted in complete filamentation (33-38→6×Pro) or a severe elongation defect (L_38_-6×Pro-P_39_; 86%).

Overall, the response observed is consistent with the AF2 model. The G_36_xxP_39_ motif is likely ideal for placing the two FtsW-binding domains of FtsL (the TM and juxtamembrane helices) in the correct orientation, though our data indicate that it is not essential. The fact that added flexibility leads to notable division defects without completely abrogating function suggests that the juxtamembrane and transmembrane binding sites cooperate in stabilizing the FtsL-FtsW interaction but are not tightly coupled structurally or functionally.

### The AlphaFold2 model corresponds well with known structural data

The good correspondence between the AF2 model and our mutational data suggests that the model is a reasonably good predictor of the structure of the FtsL-FtsW subcomplex. Although the complete structure of the FtsQLBWI complex has not been solved experimentally, the AF2 model of FtsQLBWI (illustrated in Fig. 5a) is in good agreement with the available partial structures and computational models, as well as with potential mechanisms of action both stated by and inferred from the current literature.

**Fig. 5.**
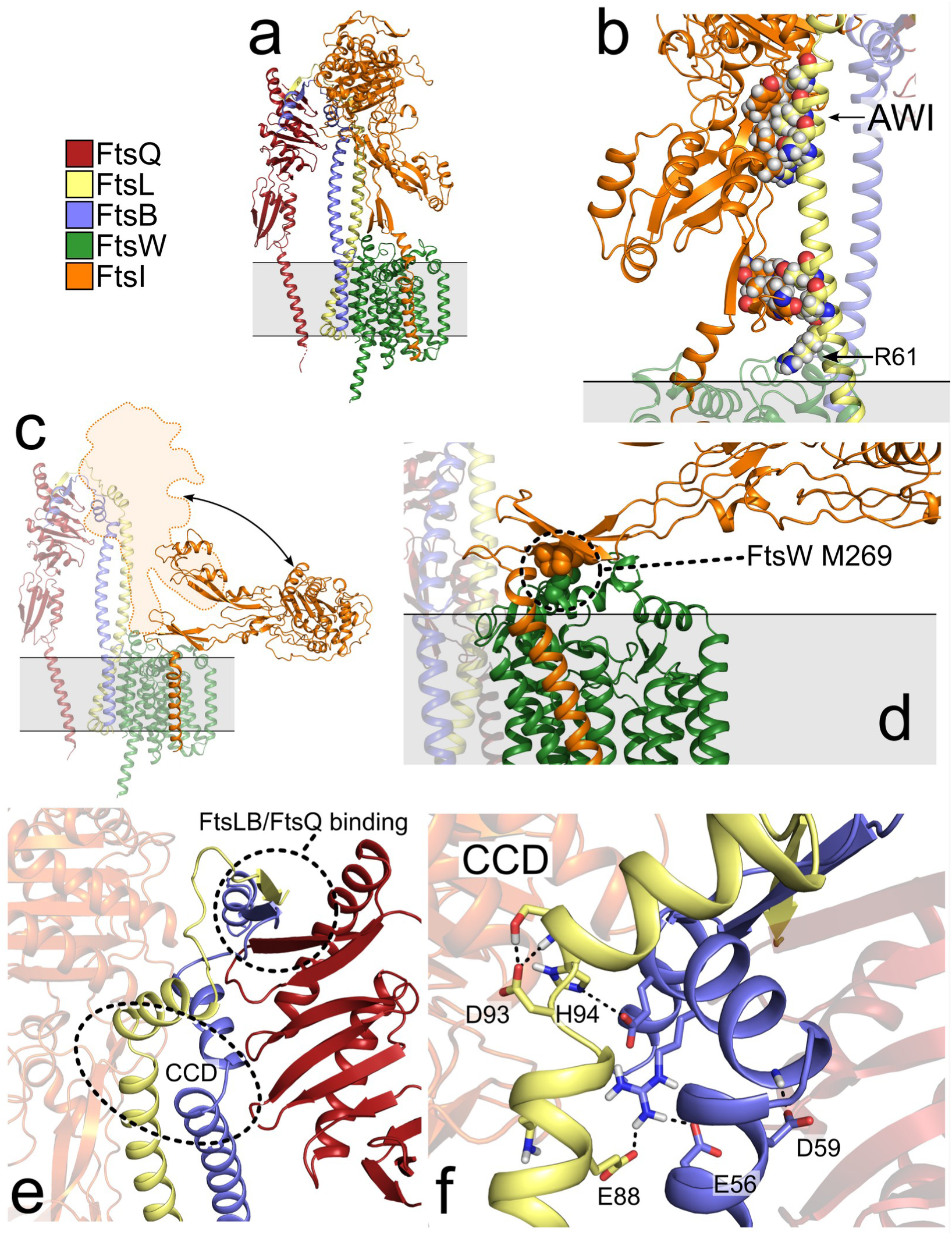
Structural organization of the AlphaFold2 FtsQLBWI model. a) Overview of the AF2 model of FtsQLBWI. b) The AF2 model captures FtsI in an active-like configuration, with its pedestal domain in contact with positions of the AWI region of the FtsL (indicated). FtsL positions R82, N83, L86, and A90 are in contact with FtsI positions V84, V86, P87, Y168, and P170. Another extensive contact occurs between the second lobe of the pedestal domain and FtsL residues R61, T64, A65, and E68, with R61 also participating in interactions with ECL4 of FtsW (positions N283 and S284). c) The AF2 model was reconfigured into a “compact” conformation using the crystal structure of RodA-PBP2 as the template. FtsI^peri^ rotates by approximately 90°, the pedestal domain moves away from FtsLB so that it is no longer in contact with the complex. d) The interaction between the FtsW ECL4 and the FtsI pedestal domain is mediated by FtsW^M269^, a superfission position, which forms a packed hydrophobic core with FtsI residues L62, V64, and I208) e) The CCD region occurs in proximity of a hinge at the end of the coiled coil. f) The majority of the CCD positions appear to stabilize the interaction with two short post-CCD helices, with residues acting as helical capping motifs (FtsL^D93^ and FtsB^D59^) or are otherwise involved in hydrogen bonding interactions. Models are available for download at the Dryad Digital Repository (DOI TBD).

First, the FtsLB component of the AF2 model is in good agreement with our recently published “Y-model” of this subcomplex, except for the oligomeric state (29). Based on experimental evidence (27,28), the Y-model was modeled as a heterotetramer (L_2_B_2_), whereas the AF2 model predicts the complex in a dimeric (L_1_B_1_) form. Despite this discrepancy, the AF2 model is nearly identical to each half of the two-fold symmetrical Y-model, lending support to the accuracy of the predictions.

AF2 also faithfully captures the interaction between FtsLB and FtsQ, which was revealed by the crystal structure of the periplasmic domain of FtsQ in complex with the C-terminal tail of FtsB (53,54). This interaction occurs through β-sheet augmentation in which the C-terminal tail of FtsB binds to the edge of the C-terminal β-sheet of FtsQ. Interestingly, the AF2 model predicts that the C-terminal tail of FtsL contributes an additional strand to the β-sheet by binding to the FtsB β-strand (Fig. 5e). This suggests that the C-terminal tail of FtsL may indirectly aid the FtsB-FtsQ interation, which is in line with prior evidence showing that a C-terminal truncation of FtsL (Δ114-121) causes cell division defects and has diminished interaction with FtsQ (34).

The AF2 model of FtsW is also in good agreement both with crystal structures of the homologous RodA (alone and in complex with the FtsI homolog PBP2) (23,45) as well as with an earlier Rosetta model of FtsW based on co-evolutionary information (50). The configuration of the TM domain of FtsI in complex with FtsW in the AF2 model is also similar to what is seen in those structures. Finally, the structure of the soluble periplasmic domain of FtsI (FtsI^peri^) closely resembles its crystal structure (17).

One notable difference between the AF2 model and the RodA-PBP2 crystal structure (23) is the orientation of the FtsI^peri^ domain. In the AF2 model, FtsI^peri^ adopts an “extended” configuration, projecting away from the membrane plane (Fig. 5a). This is in contrast to the RodA-PBP2 crystal structure, in which the soluble domain of PBP2 achieves a more “compact” configuration by lying parallel to the membrane plane, with the pedestal domain of PBP2 in contact with the RodA ECL4 (i.e., the 7/8 loop).

### The AlphaFold2 model is consistent with an activated conformation of FtsI

The compact configuration of PBP2 in the RodA-PBP2 structure (23) places its TPase catalytic site far from the PG layer, which lies approximately 120 Å above the inner membrane in Gram-negative bacteria (55). This suggests that PBP2 would be unable to crosslink novel PG stands into the existing cell wall if this were its only form. However, through negative-stain electron microscopy analysis, Sjodt *et al*. revealed that the PBP2 TPase domain can adopt a range of conformations, including extended conformations that would place it in proximity to the existing cell wall and thus in a favorable orientation for crosslinking newly synthesized PG strands (23). They hypothesized that the transition between the compact and extended configurations of the TPase domain could be critical for activation of the complex. For the elongasome-specific PBP2, this transition is likely regulated by MreC, a single pass membrane protein whose periplasmic domain binds to the pedestal domain of PBP2 (56); however, a divisome-specific homolog of MreC has not been identified. It has been proposed that this role during division may be fulfilled by the FtsL interaction with FtsI (41), and the AF2 model lends credence to this hypothesis seeing as the rigid FtsLB coiled coil appears to function as structural support for the extended orientation of FtsI^peri^. This is consistent with the prior observation that Ala insertions in the periplasmic juxtamembrane region of FtsL, which may compromise the integrity of the FtsL helix and reduce the rigidity of the coiled coil, results in a completely filamentous phenotype (28).

As further support for the hypothesis that the AF2 model captures FtsI^peri^ in an active-like configuration, it is notable that the pedestal domain of FtsI is in contact with a number of the positions in the critical AWI region of the FtsL coiled coil (Fig. 5b). Specifically, FtsL positions R_82_, N_83_, L_86_, and A_90_ are in contact with FtsI positions V_84_, V_86_, P_87_, Y_168_, and P_170_, which belong to the β-sheet connecting one of the two lobes of the pedestal domain to the TPase domain. The remaining two AWI positions, L_84_ and E_87_, do not contact FtsI in the model. Additionally, the model reveals another extensive contact between the second lobe of the pedestal domain and FtsL residues R_61_, T_64_, A_65_, and E_68_, with R_61_ also participating in hydrogen boding interactions with the backbone carbonyl groups of the extended 7/8 loop (ECL4) of FtsW (positions N_283_ and S_284_). FtsL R_61_ is another of the sensitive positions reported by Park et al. during their identification of the AWI region (41), suggesting that this interaction may have a role in orienting the ECL4 during activation.

To ask whether a compact configuration for FtsI is compatible with the AF2 model, we reconfigured FtsQLBWI in the conformation of RodA-PBP2 (23), using its crystal structure as the template for homology modeling. As schematically illustrated in Fig. 5c, FtsQLB can coexist with FtsI in a compact conformation. In the reconfiguration of the model, FtsI^peri^ is rotated by approximately 90° around a hinge that corresponds to the linker between the TM domain and the base of the pedestal domain. This moves the pedestal domain away from FtsLB so that it is no longer in contact with the complex. We conclude that FtsLB is well positioned to support FtsI^peri^ in the extended conformation and is in a proper location relative to FtsWI to allow FtsI^peri^ to adopt a wide range of motion, including the compact configuration observed in the RodA-PBP2 crystal structure. Interestingly, in the homology model, the interaction between the FtsW ECL4 and the FtsI pedestal domain appears to be mediated by FtsW^M269^, which forms a tightly packed, hydrophobic core with three FtsI residues (L_62_, V_64_, and I_208_) (Fig. 5d). This position corresponds to the known FtsW superfission mutant M269I; therefore, its involvement at this interface in the homology model supports the critical interaction between the ECL4 and the pedestal domain as a modulator of the activation mechanism for FtsWI (24). Notably, this tightly packed, hydrophobic interaction is not observed in the original structural template of RodA-PBP2, in which M_269_ corresponds to a smaller and less hydrophobic serine residue (S_224_, supplementary Fig. S8).

### Is the CCD configuration in the AlphaFold2 model in an *on* or *off* state?

In a current functional model, FtsWI activation is promoted by a transition in FtsLB from an *off* to *on* state following accumulation of FtsN at the division site. A critical component for this transition is the CCD regions of both FtsL and FtsB (35,36), which exist proximal in space and sequence to the AWI region of FtsL without overlapping it. As illustrated in Fig. 5e, the CCD occurs at the very end of the coiled coil preceding a hinge that is formed by a series of Gly residues (FtsB G_62_ and G_63_ and FtsL G_92_). The prediction of this hinge by the AF2 model is consistent with our previous observations of hinge formation obtained using molecular dynamics simulations with our Y-model (28,29). The hinge is followed by two short helices (approximately three helical turns in both FtsL and FtsB) that interact with each other in a nearly orthogonal orientation through a hydrophobic interface. The helices are then followed by unstructured, presumably flexible regions leading to their FtsQ-binding sites. The CCD region is likely critical for a rearrangement that switches FtsLB conformation from an *off* to an *on* state that is competent for activating FtsWI by enabling the interaction between FtsI^peri^ and the AWI region of FtsL. The known hyper-activated CCD mutants are likely substitutions that reduce the energy barrier of the *off/on* transition, allowing the complex to spontaneously activate, even in the absence of the FtsN signal normally necessary to overcome such a barrier.

It is interesting that a majority of the wild-type residue identities within the CCD region appear to stabilize the structure of the AF2 model. As illustrated in Fig. 5f, FtsB D_59_ and FtsL D_93_ act as capping side chains for the two short helices following the post-CCD hinge by hydrogen bonding with an unsatisfied terminal N-H backbone group. Positions FtsB E_56_ and FtsL E_88_ form salt bridges with FtsB R_70_, which belongs to the short helix. Finally, FtsL H_94_ is in close proximity to FtsB E_74_ within the short helix, though a salt bridge does not form between these two residues in the model. Notably, all known CCD hyper-activated mutations at these positions (36) either neutralize or reverse the polarity of these side chains (FtsB E56A/G/K/V, FtsB D59H/V, FtsL E88V/K, FtsL D93G, and FtsL H94Y), which would likely disrupt the stability of this region. The apparent stabilizing effect of the wild-type CCD residue identities combined with likely destabilization by the mutations suggests that the AF2 model is a good candidate for the *off* configuration of the CCD. On the other hand, FtsI binds the AWI region in an extended conformation, suggesting that the complex may instead be captured in its post-activation state. In the absence of a prediction of the other state of FtsLB (whether it is the *on* or *off* state), it is difficult to draw a mechanistic hypothesis for activation other than inferring that the hinge at the CCD region is likely to play a key role in operating the transition between the states.

### The FtsQLBWI model is compatible with a diprotomeric configuration in the “inactive” configuration

The AF2 models for both FtsLB and FtsQLBWI predict a single chain for each of the components of the complexes, forming Fts[LB]_1_ and Fts[QLBWI]_1_ monoprotomers, respectively. These stoichiometries are in conflict with our previous *in vitro* evidence which supports two chains of both FtsL and FtsB forming an Fts[LB]_2_ heterotetramer (27,28). As discussed previously, the Fts[LB]_1_ AF2 model is very similar to half of our Fts[LB]_2_ Y-model, which raises the possibility that AF2 may be missing the key dimerization interaction between the FtsLB protomers in the transmembrane region. If this interaction were maintained during divisome assembly, FtsLB could therefore act as a central hub, binding two FtsWI complexes as well as two FtsQ subunits into a single complex. For this reason, we tested if the AF2 model is compatible with a hypothetical Fts[QLBWI]_2_ diprotomeric configuration. This was performed by applying the same C2 symmetry operation (180° rotation) to the Fts[QLBWI]_1_ complex (Fig. 6b) that was used to reconstruct the Fts[LB]_2_ Y-model from the Fts[LB]_1_ AF2 prediction (Fig. 6a).

**Fig. 6.**
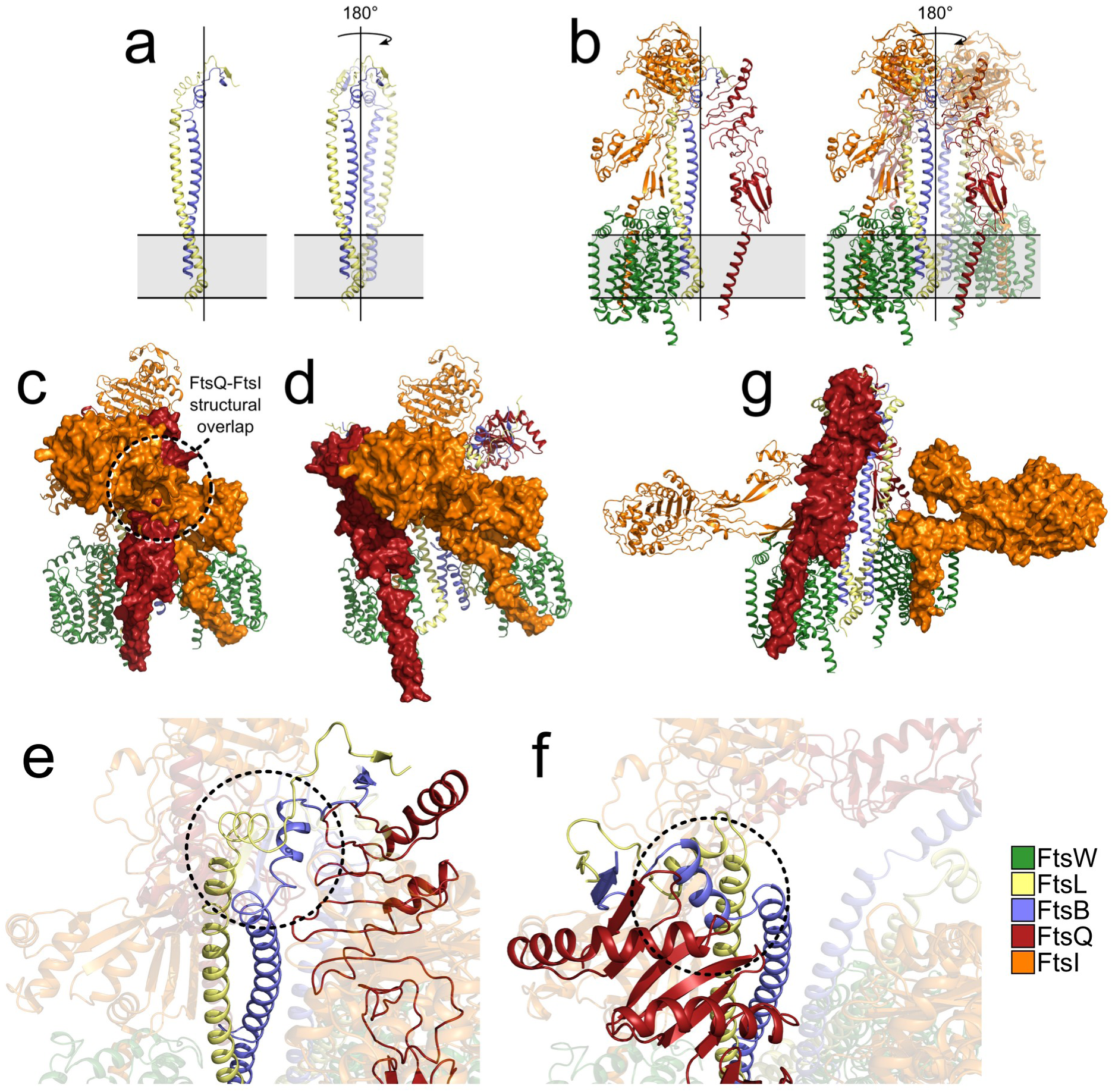
The FtsQLBWI complex in a hypothetical diprotomeric configuration. a) The diprotomeric configuration of FtsLB (Y-model) can be obtained by applying two-fold symmetry along the main axis of the helical bundle. b) The same symmetry operation can be applied to the AlphaFold2 model to obtain a diprotomeric configuration for the entire FtsQLBWI complex. c) In this diprotomeric configuration, FtsQ ^peri^ and FtsI^peri^ in extended conformation overlap in space. d) FtsQ can be relocated in a position that is compatible with FtsI by applying flexibility to the hinge in the CCD region, as illustrated from above in (e) the original clashing configuration and in (f) the reconfigured hinge and FtsQ location. g) The diprotomeric complex is compatible with the hypothetical compact state of FtsI.

FtsQLBWI appears compatible with a diprotomeric configuration except for two sets of steric overlaps (one minor and one severe), both of which involve the FtsQ subunits. The first issue is an overlap between the terminal regions of FtsQ and FtsLB from opposite protomers. FtsQ binds to the terminal tails of FtsLB following a linker region that is presumably quite flexible (54), but AF2 unfortunately places the FtsQ subunit in the same conflicting spatial orientation in all five solutions. However, when the FtsQLB ternary complex is predicted by AF2, the resulting models display a wider variability in the spatial orientation of FtsQ with respect to FtsLB (supplementary Fig. S9). We found that an orientation corresponding to rank model #4 of FtsQLB is compatible with a diprotomeric configuration when introduced into Fts[QLBWI]_2_.

A more severe steric overlap occurs between FtsQ^peri^ and FtsI^peri^, which occupy the same region of space adjacent to FtsLB in the Fts[QLBWI]_2_ model (highlighted by a circle in Fig. 6c). Unlike the steric clash with the FtsQLB C-terminal tails, this overlap cannot be resolved with the above procedure. To address whether this clash could be solved by providing flexibility to the CCD hinge we used a different procedure, based on docking of FtsQ^peri^ with HADDOCK followed by loop reconstruction with Rosetta, as detailed in the Methods. We found that with this procedure FtsQ can be relocated in an orientation that no longer overlaps with FtsI^peri^ in its extended configuration (Fig. 6d). The rearrangement of the CCD hinge is illustrated in detail in Fig. 6e,f.

Notably, the FtsQ/FtsI overlap can be completely resolved also by reconfiguring Fts[QLBWI]_2_ with FtsI in the “compact” state, since this configuration moves FtsI ^peri^ away from the central FtsLB hub (Fig. 6g). This finding is interesting because it places FtsQ as a potential gatekeeper of the interaction between the FtsL AWI region and FtsI^peri^. This function would be dependent on the orientation of FtsQ, which in turn is determined by a conformational change of the FtsLB CCD hinge. According to this hypothesis, we provide a speculative model for FtsQLB-dependent FtsWI activation in the Conclusions section of this article.

## Conclusions

In this article, we investigate the essential interaction between FtsL and FtsW, which is necessary for FtsW to be recruited to the division site. We identified mutations in the juxtamembrane and transmembrane regions of the two proteins that result in division defects and loss of localization of FtsW. These mutations are in overall good agreement with the predicted interface between FtsL and FtsW obtained using AF2 modeling. This model identifies two major FtsL-FtsW interfaces – a cytoplasmic interaction between a short, juxtamembrane helix in FtsL and the proximal regions to TM1 and TM10 in FtsW as well as a transmembrane interaction between the TM helix of FtsL and TM10 of FtsW. We also tested different structural requirements of the linker region between these two interfaces in FtsL. We found that modifying length and flexibility of this linker can lead to division defects, suggesting that the two interaction sites cooperate in stabilizing the interaction that mediates the formation of FtsQLB and FtsWI into a complex.

Given the good correspondence between the AF2 model and current understanding of the FtsQLB and FtsWI complexes, we used computational tools to remodel the structure, seeking insight into the hypothesized structural transitions that first switch FtsLB from an *off* to an *on* state and then lead to the activation of the FtsWI complex and PG synthesis. Overall, the original AF2 model suggests that FtsLB serves as a support for FtsI^peri^ in its extended conformation (41), orienting the TPase domain towards the PG layer through interactions with the FtsL AWI region. The structural model also appears compatible with a rearrangement that puts FtsI^peri^ into a “compact” conformation that lays on top of the cytoplasmic face of FtsW, parallel to the membrane. In this potential transition between the compact (presumably inactive) state and the extended (presumably active) state, changes occur in the contacts between the pedestal domain of FtsI and the critical 7/8 loop (ECL4) of FtsW, which could lead to derepression of FtsW GTase activity (24).

The AF2 model provides a useful structural hypothesis for the activation of the synthase complex, but the nature of the *off*/*on* transition in FtsLB that promotes activation is less clear. The critical element for this transition is the CCD, but the model does not offer a clear indication of its alternative states. Since the CCD region occurs in correspondence with a hinge, a conformational change around this hinge would most likely induce a shift in the position of FtsQ with respect of the rest of the complex. It is thus possible that the *off* state is one in which FtsQ is positioned in a way that shields the FtsL AWI region from interactions with FtsI. In the monoprotomeric AF2 model, FtsQ would have to move to the opposite side of FtsLB to shield the AWI positions, which is a possible mechanism. In the hypothetical diprotomeric configuration of Fts[QLBWI]_2_, the region around the central FtsLB hub is crowded, and FtsQ from the opposing protomer is placed in a conformation that would prevent binding of FtsI ^peri^ to the FtsL AWI region.

Based on the above considerations, we offer a speculative model for regulation and activation of the FtsWI sythase complex, illustrated in Fig. 7. We propose that the FtsQLBWI complex (potentially in a diprotomeric configuration) travels along with the treadmilling FtsZ protofilaments in its inactive state (panel *a*). The SPOR domain of FtsN recognizes regions of the cell wall that are enriched in denuded PG (glycan chains devoid of peptide stems), signaling a site in which new material can be inserted (38,39). Interaction with FtsN could result in a structural reconfiguration of Fts[QLBWI]_2_ that shifts FtsQ into a position that no longer shields the FtsL AWI region (panel *b*). The structural reconfiguration may be promoted in part by the tension that arises between the moving Fts[QLBWI]_2_ complex (which travels with the treadmilling FtsZ filaments) and the stationary FtsN bound to septal PG.

**Fig. 7.**
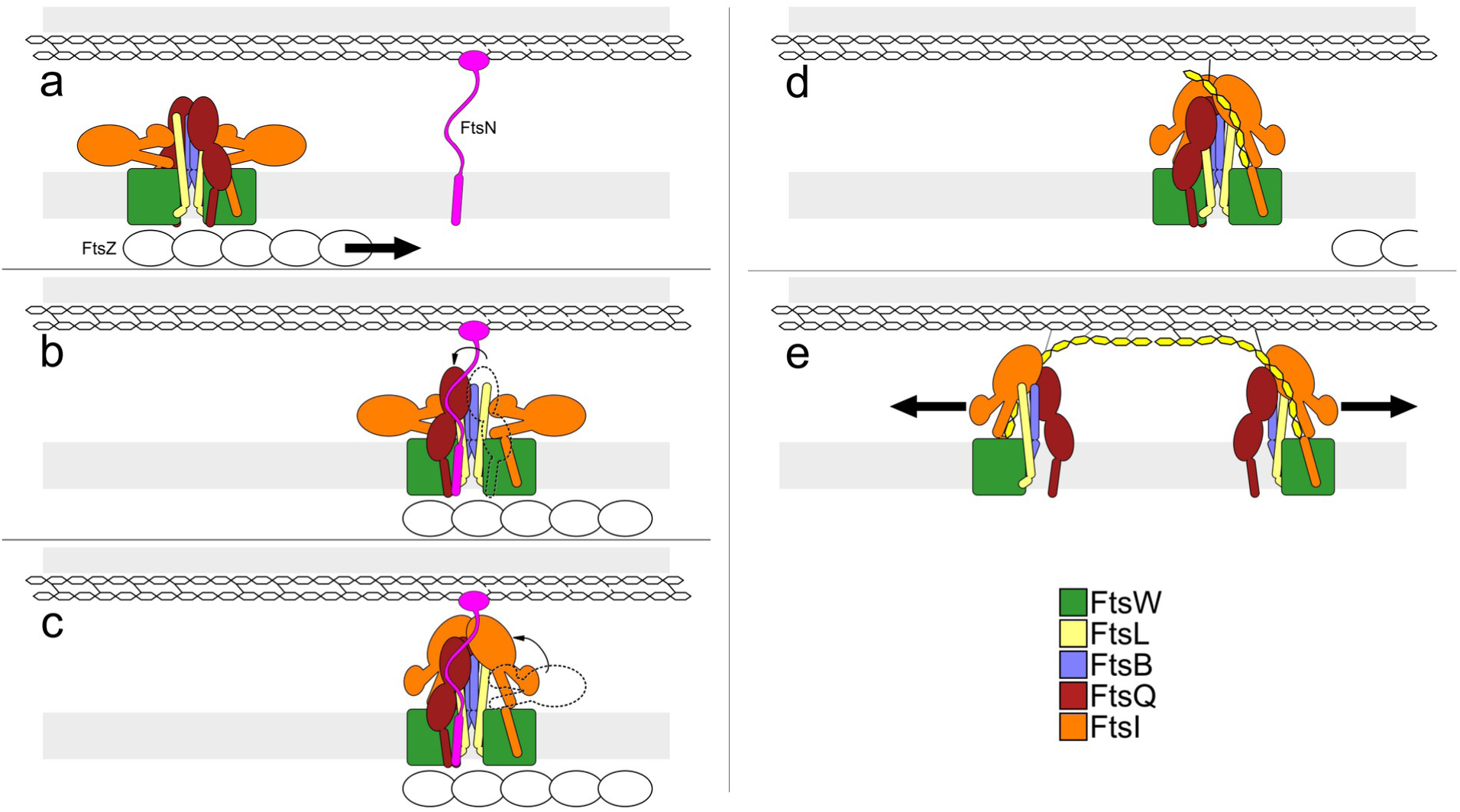
A proposed hypothetical mechanism for septal PG synthesis activation based on a diprotomeric configuration of FtsQLBWI. a) The Fts[QLBWI]2 complex travels with the treadmilling FtsZ filaments in a hypothetical diprotomeric state. b) The complex reaches a region of denuded PG where FtsN has accumulated. Binding to FtsN reconfigures the position of FtsQ, exposing the AWI region of FtsL. c) Unhindered by FtsQ, FtsI can assume an extended conformation, binding to the AWI region and activating FtsW. d) Fts[QLBWI]2 is released from FtsZ and PG synthesis begins. e) The mechanism does not necessarily imply a diprotomeric complex, but if that occurs, the two synthase complexes may start depositing PG with opposite polarities. We speculate that it may be possible for the diprotomeric complex to split into two independently working protomers.

The reconfiguration of FtsQ would then enable the transition of FtsI^peri^ from the compact to the extended conformation, supported by the coiled-coil domain of FtsLB (panel *c*). In this orientation, FtsI activates FtsW synthesis through interaction between residues in the pedestal domain and the ECL4 of FtsW, and the complex is subsequently released from FtsZ (9) (panel *d*). In the final step of our speculative model, we speculate on the possibility that a diprotomeric complex would split into two independently functioning protomers, based on the fact that the two protomers are oriented in diametrically opposed directions, which may lead to deposition of PG strands with opposed polarity. The shear forces generated by synthesis could potentially lead to separation of the protomers. Therefore, the deposition of the newly synthesized PG would have the same origin, but the strands would elongate in opposing directions (panel *e*).

Although the model is speculative, it is in reasonable agreement with the current views about the complex and its regulation. In addition, it has the virtue of providing a possible mechanistic role for the CCD, which is still unknown. Recently, Attaibi and den Blaauwen proposed a different model for the control of septal PG synthesis also based on a higher oligomeric state for the complex (49). Their model also considers FtsK, which exists in a hexameric state and binds to FtsQ (57–59). Accordingly, they suggested that FtsQLBWI exists in a hexaprotomeric state Fts[KQLBWI]_6_. In their model, the complex would remain in this higher oligomer state after activation and would thus synthesize a six-stranded layer of PG with the same polarity. The authors propose that this type of deposition could lead to the invagination of the cell wall.

Whether these hypothetical higher-order complexes remain associated (working together as proposed by den Blaauwen) or rather separate (in line with our model) is an open question. Experiments to determine the dynamics of the complex *in vivo* and the oligomeric state of FtsQLBWI in its active and inactive states are necessary to test these models and other potential alternatives. In addition, further work is required to dissect the fine molecular mechanisms that control the activation of PG synthesis and reconstruction of cellular shape during cell division.

## Materials and methods

### Plasmid cloning

For the *in vivo* complementation experiments, mutant variants of FtsL or FtsW were cloned via standard QuikChange mutagenesis or inverse PCR into pMDG29 (34) (FLAG3-FtsL) or pSJC214 (GFP-FtsW), respectively. All other constructs were amplified from various lab plasmids. Further plasmid editing was performed via standard Gibson assembly (60) and QuikChange mutagenesis. All constructs were confirmed by DNA sequencing (Quintara Biosciences or Functional Biosciences). A complete plasmid inventory is included in supplementary Table S2.

### Bacterial strains and media for *in vivo* experiments

The phenotypic analyses were performed using depletion strains MDG277 (33) or MDG254 (34) for FtsL and EC912 (18) for FtsW. For all experiments described, bacterial cells were grown in LB medium supplemented with 100 μg/mL spectinomycin (Dot Scientific) or 100 μg/mL ampicillin (Thermo Fisher) and the appropriate carbon source. Medium was supplemented with 0.2% (w/v) L-arabinose (Sigma) or 0.2% (w/v) D-glucose (Sigma) to induce or repress, respectively, the expression of the WT genes regulated by a P_BAD_ promoter. 20 μM IPTG was added to the media to induce the expression of genes regulated by a P_trc_ promoter on the transformed plasmid.

### Depletion strain experiments

The protocol for the depletion strain experiments was adapted from prior work (18,33). In short, a mutated copy of FtsL or FtsW was transformed into its respective depletion strain. Strains were grown overnight at 37 °C on an LB plate supplemented with arabinose and appropriate antibiotics. A single colony from the plate was grown overnight at 37 °C in 3 mL of LB medium supplemented with arabinose and antibiotic. The overnight culture was then diluted 1:100 into fresh LB medium containing the same supplements and grown to an OD_600_ of ~0.3. An aliquot of 1 mL of culture was washed twice with LB medium lacking any sugar and then diluted 1:100 into 3 mL of fresh LB medium supplemented with glucose, IPTG, and antibiotic to induce expression of the mutated gene and to repress the WT gene. The cells were then grown at 37 °C for 3.5 h (MDG277 and MDG254) or 4.5 h (EC912), which is the approximate time necessary to deplete the cells of the WT protein. The cells were then placed on ice to stop growth before imaging. Depletion strains were provided with a copy of their respective WT protein in the plasmid for a positive control. Similarly, depletion strains with no protein in the plasmid (empty vector) were tested as negative controls.

### Microscopy and cell length measurements

10 μl of cell samples were mounted on a number 1.5, 24 × 50 mm (0.16 – 0.19 mm thickness) cover glass slide (Thermo Fisher or VWR). Cells were cushioned with a 3% (w/v) agarose gel pad to restrict the movement of the live cells. Cells were optically imaged using a Nikon Eclipse Ti inverted microscope equipped with crossed polarizers and a Photometrics CoolSNAP HQ2 CCD camera using a Nikon X100 oil objective lens. Phase-contrast images of bacterial cells were recorded with a 50 ms exposure time using Nikon NIS Elements software. GFP images were recorded with a 3-5 s exposure time. Multiple snapshots were collected for each experiment. For FtsL, all images were analyzed to measure the cell length in Oufti (61) using one single optimized parameter set and manual verification, as previously described (28). Cells with lengths longer than the 95% percentile in the WT length distribution were considered “elongated,” and length distributions for the various mutations were compared to this threshold. In general, we refer to phenotypes with <10% of total cells being elongated as “WT-like” or “no defect,” ≥10% as “mild,” ≥25% as “moderate,” ≥50% as “severe,” and “filamentous” when complete filamentation occurred. Completely filamentous mutations resulted in a low cell count which generally precluded cell length distribution analysis.

For FtsL and FtsW, images were additionally analyzed using Misic, a deep learning method to segment microscopy images of rod-shaped cells (62). Cell images were segmented with the noise parameter set to 0.13 and the cell width set to 20 pixels. The watershed method was used for post-processing, and for each image an ordinal mask was generated where the pixels corresponding to each individual cell were all given a unique intensity value. This mask was processed using MicrobeJ (63) to create cell contours and measure the cell lengths. When necessary, cell contours were manually corrected to fix egregious errors in the segmentation. Cells touching the image edge, as well as cell contours less than 8 or greater than 30 pixels in width, as well as contours with more than two poles were excluded from the analysis. Because Misic allowed for much greater throughput cell segmentation, cells for each mutant were classified as “elongated” based on the 95% percentile of WT cell images collected on the same dates instead of using a representative WT dataset as was done for the Oufti analysis.

### Western blots

Expression level across variants was assessed by Western blot analysis (Fig. S3). About 3 ml of cells were pelleted and resuspended in 300 μl of lysis buffer (50 mM Hepes, pH 8.0, and 50 mM NaCl) with 5 mM β-mercaptoethanol (βME). The cells were sonicated and centrifuged at 21,000g for 10 min before collecting the supernatant. Total protein concentration was determined by bicinchoninic acid assay (Pierce). 120 μl of lysates were mixed with 40 μl of 4× LDS sample buffer (Novex, Life Technologies) with βME and boiled at 98 °C for 3 min (FtsW samples were not boiled to facilitate antibody binding to GFP). For each FtsL and FtsW sample, the equivalent of 7 μg or >35 μg, respectively, of total protein was separated by SDS-PAGE (Invitrogen) and transferred to polyvinylidene difluoride membrane (VWR). For immunoblotting, horseradish peroxidase (HRP) conjugated α-FLAG (M2) antibodies (Sigma; 1:1000) were used for FtsL, whereas HRP conjugated α-GFP antibodies (ThermoFisher; 1:7000) were used for FtsW.

### Prediction of FtsBLQWI complex and subcomplexes using AlphaFold2

The structures of the *E. coli* FtsBLQWI complex and the subcomplexes were predicted with AlphaFold2 using localcolabfold, an extension of ColabFold that can be run using local computing resources (64). The full sequences of FtsQ, FtsL, FtsB, FtsW, and FtsI were used as input for multimer predictions. For the FtsLBWI and FtsQLBWI complexes, the AlphaFold2-multimer-v2 model was used, while for the FtsQLB complex, the AlphaFold2-multimer model was used (47). The Mmseqs2 UniRef+Environmental database was used to search for homologs (65). Interolog pairing was accomplished with the unpaired+paired option, which identifies sequences for the multiple sequence alignment (MSA) by genomic distance when possible. 5 prediction cycles were used, and 5 models for each prediction were generated and ranked by the ptmscore.

### Modeling the FtsQLBWI complex in the compact configuration

A homology model of FtsW-FtsI in which FtsI is oriented in a compact configuration was generated using the SWISS-MODEL webserver (66) using the RodA-PBP2 structure (PDB 6PL5) as a template. This model was aligned to the AlphaFold2 FtsQLBWI prediction, and the FtsI of the AlphaFold2 model was replaced with the FtsI from the homology model.

### Modeling a diprotomeric FtsLBWI complex

To identify a conformation of the previously-generated FtsLB Y-model (29) that was most compatible with the conformation of the AlphaFold2 FtsLBWI complex, we first created C2-symmetric averages of snapshots of the molecular dynamics simulations in R. Each snapshot was first aligned parallel to the Z-axis (membrane normal), then duplicated and rotated by 180°. Backbone coordinates were averaged using the bio3d package (67) in R, and the FtsBLWI model was aligned to these snapshots using FtsB^1-61^ and FtsL^40-91^. The snapshot with the lowest RMSD to the FtsLBWI model was selected as the basis of the diprotomer, which was generated via duplication and alignment to the opposite set of chains.

### Integrative modeling of a diprotomeric FtsQLBWI complex in the “active” conformation

We first attempted to create a C2-symmetric model of the AlphaFold2 FtsQLBWI protomer via simple rotation, but this led to extreme clashes between FtsQ and FtsI, FtsL, and FtsB on the opposite protomer. To identify whether FtsQ could be accommodated in the diprotomeric arrangement at all, we used HADDOCK 2.4 (68,69) to dock the FtsLBWI diprotomer to two segments composed of the periplasmic domain of FtsQ in complex with the C-terminal tails of FtsL and FtsB. Before the procedure, the FtsLBWI diprotomer generated from the AlphaFold2 FtsLBWI prediction was trimmed to remove low-confidence residues at the termini of each chain as well as the C-termini of FtsB and FtsL. The FtsQLB body was prepared by extracting residues FtsQ^58-257^, FtsL^95-12^, and FtsB^64-88^ from the AlphaFold2 FtsQLBWI prediction. The hinge regions of FtsL and FtsB, corresponding to FtsL^92-94^ and FtsB^61-63^ were removed from all segments.

The transmembrane domains of the FtsLBWI diprotomer were kept fixed during the rigid-body docking step, while the FtsQLB segments were free to move. Connectivity between the FtsL and FtsB chains was preserved via the addition of unambiguous distance restraints up to 12 Å between the alpha carbons of FtsL^91^:FtsL^95^ and FtsB^60^:FtsB^64^. Symmetry restraints were added to each matching chain in the FtsLBWI diprotomer segment as well as the matching chains in the FtsQLB segments. Additionally, to preserve global C2-symmetry, symmetry restraints were added to the unambig.tbl restraint file such that the distance between each alpha carbon in one FtsQLB segment to FtsB^10^, FtsB^41^, and FtsB^56^ from one FtsB chain matched the distance of the other FtsQLB segment to the same residues from the other FtsB chain. To ensure that the docked results were compatible with the FtsQ transmembrane domain, harmonic Z-distance restraints were added to FtsQ^S58^ on each segment such that they would be kept between 20-40 Å above the origin (center of the membrane). 4,000 models were generated during the rigid-body docking phase, and the best 400 were selected for semi-flexible annealing and refinement.

#### Remodeling the hinge region of FtsL and FtsB

Candidate orientations of the FtsQLBWI complex were filtered for proper connectivity of FtsL and FtsB by attempting to rebuild the hinge regions of FtsL and FtsB using the remodel application in Rosetta (70). The FtsL and FtsB of one protomer were extracted from the docked complexes and used to create a blueprint file. The missing hinge residues were added to the blueprint and allowed to adopt a random secondary structure. One residue on either side of the hinge region was allowed to be flexible and to take a random secondary structure. The quick_and_dirty protocol was used, and 100 iterations were attempted for each candidate. Candidates with no successfully rebuilt hinge regions were excluded from further analysis.

#### Remodeling the FtsQ TM helix and juxtamembrane linker

Candidate orientations of the FtsQLBWI complex were additionally filtered for proper orientation of FtsQ by alignment with alternative FtsQ conformations (described below) via a constrained alignment procedure. First, for each alternative conformation, the X and Y coordinates of the geometric center of the mobile selection (corresponding to the backbone atoms of the first β-strand of the FtsQ POTRA domain, FtsQ^58-63^) were translated to that of the target, keeping the Z-coordinate fixed. Next, the quaternion method was used to calculate the optimal rotation matrix to align the two selections (71). From this matrix, only the rotation about the Z-axis was performed so that the orientation of FtsQ with respect to the membrane plane was preserved. Models that aligned with an RMSD < 1 Å for the selected segment were identified, and of these, the alternative FtsQ conformation with the lowest Rosetta score that did not clash with the rest of the FtsQLBWI diprotomeric model was selected for that candidate.

#### Final merging and refinement of the FtsQLBWI diprotomer

Candidate orientations of the FtsQLBWI diprotomeric complex that were compatible with the hinge region and the FtsQ TM helix were clustered via an RMSD threshold of 7.5 Å. One model from each cluster was selected based on the haddock score and the rosetta score of the rebuilt hinge region. The hinge regions of FtsB and FtsL and the FtsQ TM helix + juxtamembrane domain were extracted from the corresponding pdb files using pdb_tools (72) and merged with each FtsQLBWI protomer via a structural alignment followed by a weighted averaging of the shared backbone coordinates using a program written in the Molecular Software Library (73) to minimize local disruptions of the structure. Finally, rotamers were repacked and the models were relaxed using the relax application of Rosetta (74) with the following flags: -relax:fast, -relax:constrain_relax_to_start_coords, -relax:ramp_constraints false, -ex1, -ex2, -use_input_sc, -no_his_his_pairE, -no_optH false, and -flip_HNQ.

### Modeling FtsQ TM and juxtamembrane linker orientations

To explore possible alternate conformations of the FtsQ TM helix and juxtamembrane linker, we used the mp_domain_assembly application in Rosetta (75). The transmembrane domain of FtsQ was generated using the helix_from_sequence application (76) in RosettaMP (77) and oriented in the membrane with the PPM webserver (78). The FtsQ chain of the crystal structure of FtsQ in complex with FtsB (PDB 6H9N) was used as the periplasmic domain. To ensure sufficient exploration of the conformational space, 100,000 models were generated.

### Modeling of a FtsQLBWI diprotomeric complex in the “inactive” state

To model the Fts[QLBWI]_2_ diprotomer in the inactive state, we first attempted to make the model of the FtsQLBWI extended protomer C2-symmetric as described above. However, this resulted in severe clashes between FtsQ and the coiled-coil domains of the FtsLB subunits in the other protomer. We next removed FtsQ from this diprotomer and aligned it to the 5 models of FtsQLB from AlphaFold2, as we noticed the relative orientations of FtsQ and FtsLB were quite diverse for these predicted structures. The 4^th^ model produced a clash-free alignment of the FtsQ periplasmic domain with the rest of the FtsLBWI diprotomer, but the TM domain clashed heavily with FtsW. We opted to find an alternate arrangement of the FtsQ TM helix and juxtamembrane linker using the alternative conformations of FtsQ and membrane-constrained alignment strategy described above. Models that aligned with an RMSD < 1 Å were identified, and of these, the FtsQ TM helix model with the lowest Rosetta score that did not clash with the rest of the FtsQLBWI structure was selected for that candidate. The FtsQ TM helix and the FtsQLB model were merged with the FtsLBWI diprotomer to produce the final arrangement, which was then relaxed using the procedure described above.

### Statistical analysis and data visualization

Statistical analysis and visualization of cell length distributions were performed in R with the aid of the following packages: tidyverse (79), magick (80), imageR (81), ggdist (82), egg (83), and bactMAP (84). Visualization of the PAE and identification of contacting residues in the AlphaFold2 predictions were performed in R with additional aid from the rjson (85) and bio3d (67) packages.

### Availability of molecular models

Five PDB files were deposited to the Dryad Digital Repository and are free to download (submission currently under processing). These include: 1) the original AF2 protomeric model; 2) the protomeric in which FtsI assumes a compact configuration; 3) the initial diprotomeric model in which the periplasmic domains of FtsQ and FtsI clash; 4) the reconfigured diprotomeric model in the extended conformation; 5) the diprotomeric model in the compact configuration.

## Acknowledgments

This work was supported in part by National Institutes of Health (NIH) Grant R35-GM130339 to A.S. This research was performed using the computing resources and assistance of the UW-Madison Center For High Throughput Computing (CHTC) in the Department of Computer Sciences. We would also like to thank the Jon Beckwith and David Weiss labs for providing strains and plasmids used in this work.

## SUPPLEMENTARY INFORMATION

### Supplementary Tables

**Table S1.** Summary of the phenotypes of mutations made within the cytoplasmic face of FtsW. The elongation percentage indicates fraction of total cell population that is elongated. Elongated cells are defined as those exceeding the length of the 95^th^ percentile in the WT length distribution. Localization: “++” refers to WT-like frequency of GFP foci; “+” refers to a small reduction in GFP foci frequency; “-” refers to a major decrease in GFP foci frequency; “--” refers to the complete or near absence of GFP foci at division sites (categorization was performed by eye). The representative images are included in supplementary Fig. S5. Cell length distrubutions are plotted in supplementary Fig. S4. FtsW mutations are also summarized in Fig. 3.

| **FtsW mutation** | **Location in FtsW** | **% elongation** | **FtsW localization** |
| --- | --- | --- | --- |
| + control (WT) | NA | 5 | ++ |
| - control (EV) | NA | 100 | - - |
| $\Delta 1-42$ | N-terminus | 7 | ++ |
| $\Delta 1-46$ | N-terminus | 84 | - |
| 32-37→6×Ala | N-terminus | 5 | ++ |
| 38-42→5×Ala | N-terminus | 3 | ++ |
| 43-44→2×Ala | N-terminus | 9 | ++ |
| 43-46→4×Ala | N-terminus | 92 | - |
| Y44F/R46A | N-terminus | 5 | ++ |
| Y44F/R46E | N-terminus | 7 | ++ |
| Y44A/R46A | N-terminus | 13 | ++ |
| Y44A/R46E | N-terminus | 17 | ++ |
| 45-46→2×Ala | N-terminus | 29 | + |
| R46E | N-terminus | 11 | ++ |
| F53A/A57L/F60A/I61A | TM1 | 54 | ++ |
| 105-109→5×Ala | 2/3 loop | 10 | ++ |
| R109E | 2/3 loop | 22 | + |
| 163-165→3×Ala | 4/5 loop | 76 | ++ |
| 163-168→6×Ala | 4/5 loop | 91 | ++ |
| R166E/K167E | 4/5 loop | 93 | ++ |
| 166-167→2×Ala | 4/5 loop | 89 | ++ |
| 169-171→3×Ala | 4/5 loop | 73 | ++ |
| R172E/R176E | 4/5 loop | 36 | ++ |
| 172-176→5×Ala | 4/5 loop | 48 | ++ |
| K218E | 6/7 loop | 10 | ++ |
| 325-330→6×Ala | 8/9 loop | 24 | + |
| R325E/R331E/K332E | 8/9 loop | 9 | + |
| 331-336→6×Ala | 8/9 loop | 6 | ++ |
| 337-339→3×Ala | 8/9 loop | 5 | ++ |
| 337-342→6×Ala | 8/9 loop | 78 | - |
| R339E | 8/9 loop | 7 | ++ |
| 340-342→3×Ala | 8/9 loop | 2 | + |
| 395-397→3×Ala | C-terminus | 10 | + |
| R396E | C-terminus | 10 | ++ |
| 396-401→6×Ala | C-terminus | 52 | - |
| R396X | C-terminus | 91 | - - |
| 398-399→2×Ala | C-terminus | 12 | ++ |
| 398-401→4×Ala | C-terminus | 79 | - |
| 400-401→2×Ala | C-terminus | 6 | ++ |
| R402X | C-terminus | 13 | ++ |

**Table S2.**
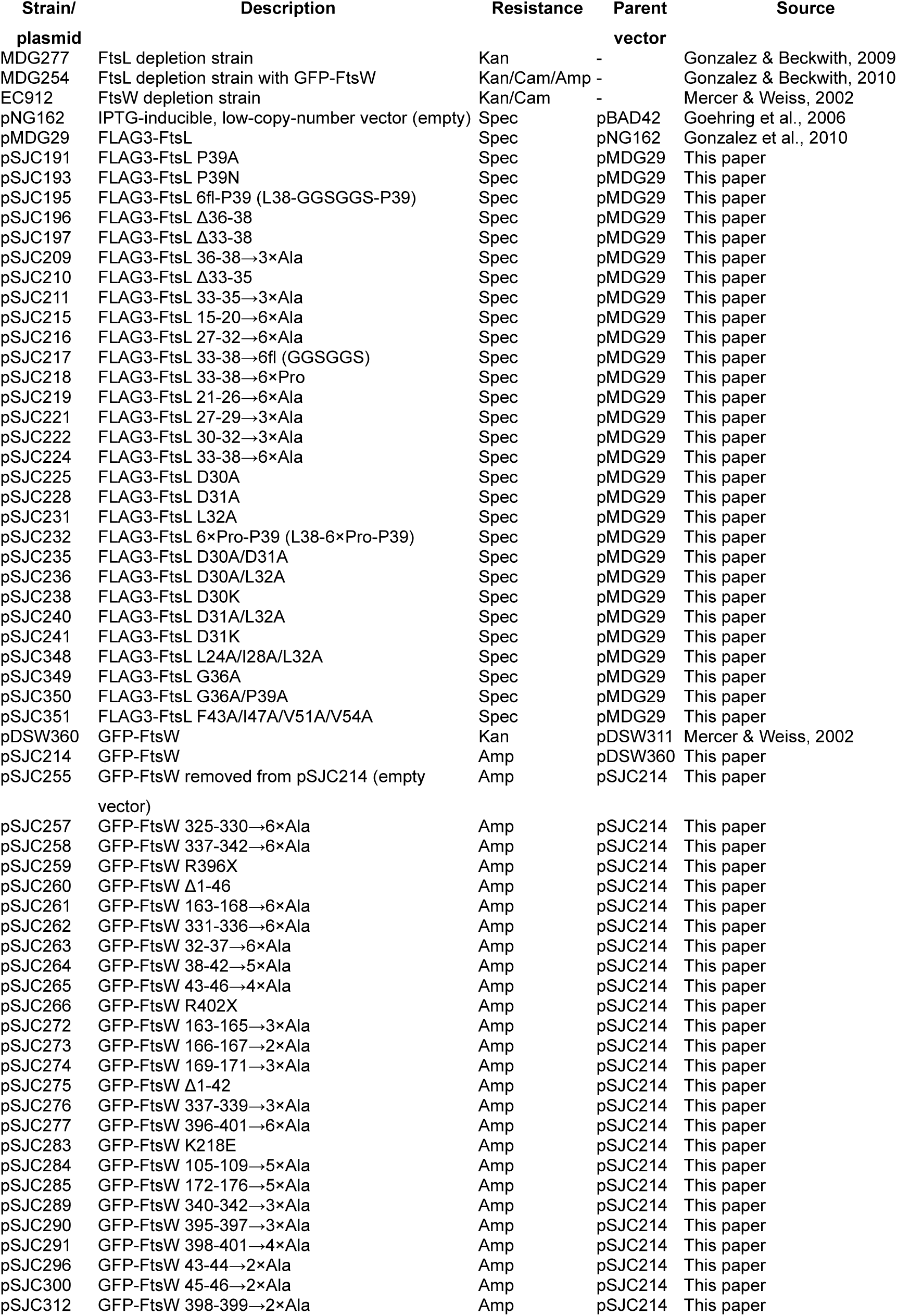
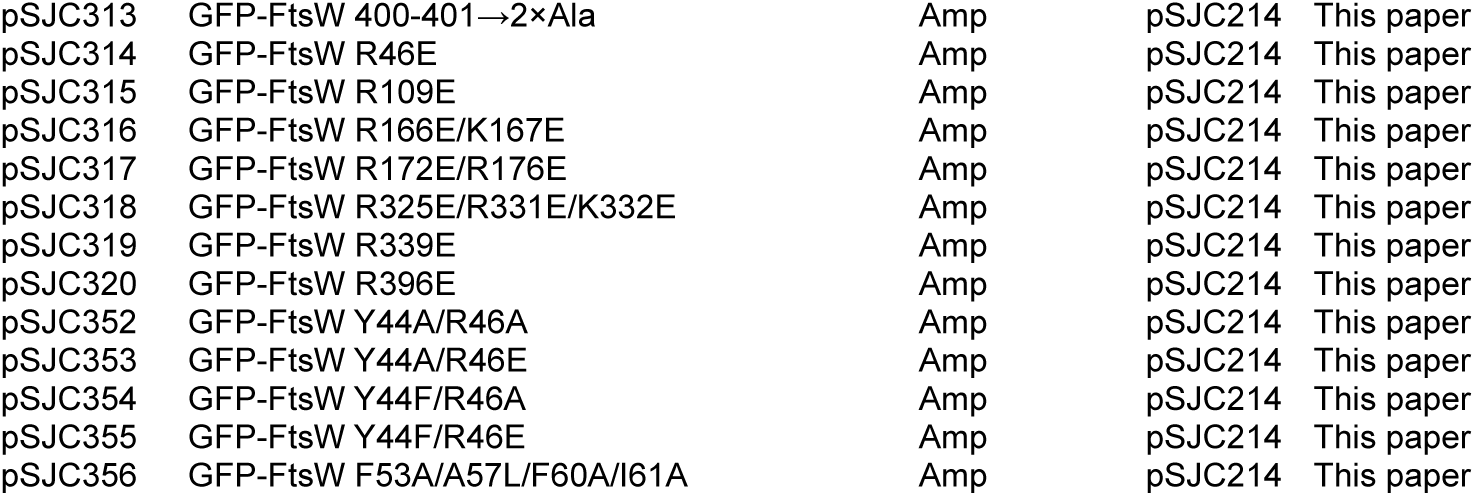
Strains and plasmids used in this paper. “X” refers to mutation to a stop codon.

### Supplementary Figures

**Fig. S1.**
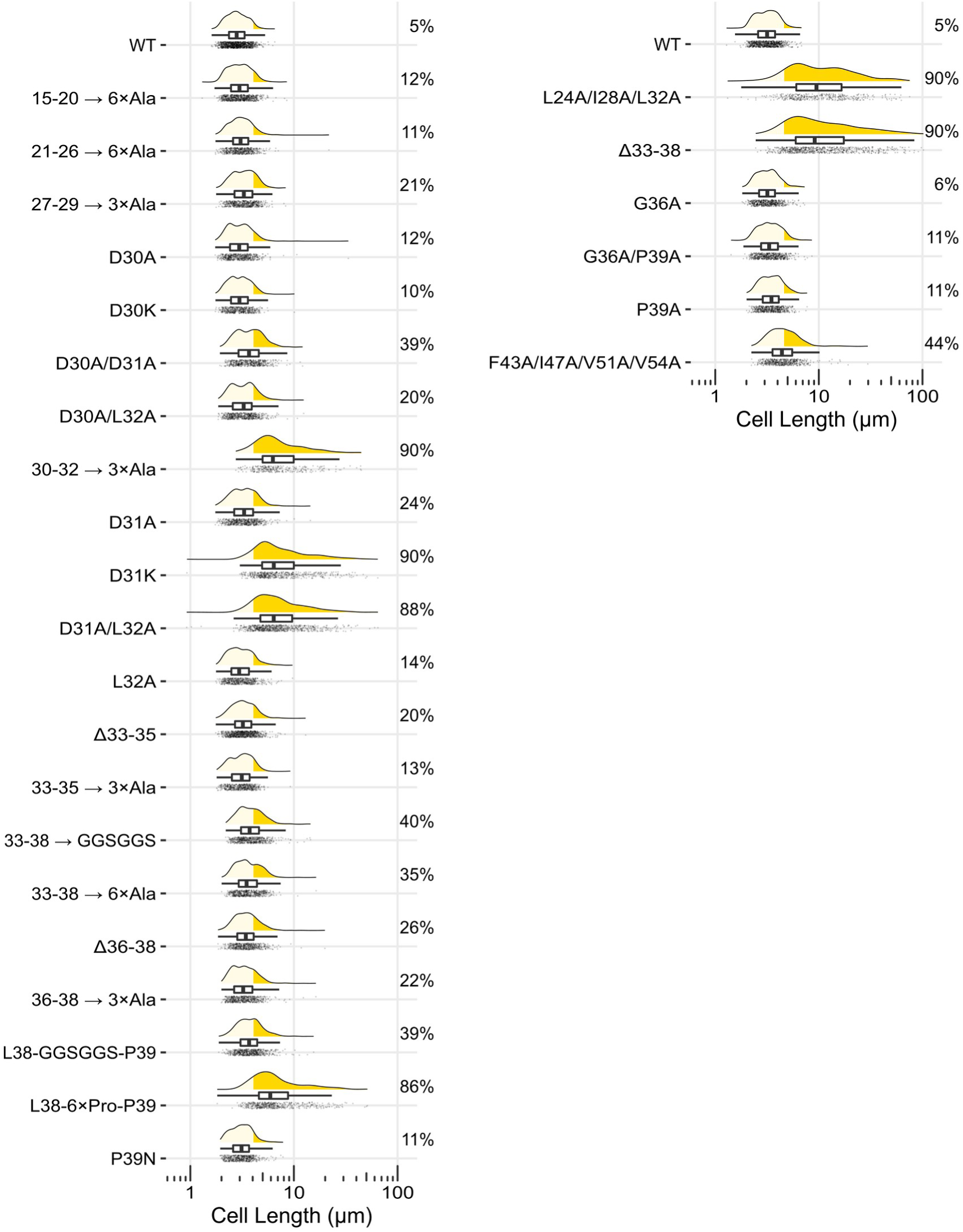
Effect of FtsL cytoplasmic tail and TM mutations. Summary of elongation defects for FtsL mutations tested within this paper. Elongated cells are defined as those exceeding the length of the 95^th^ percentile in the WT length distribution (yellow area; percentage indicates fraction of total population that is elongated).

**Fig. S2.**
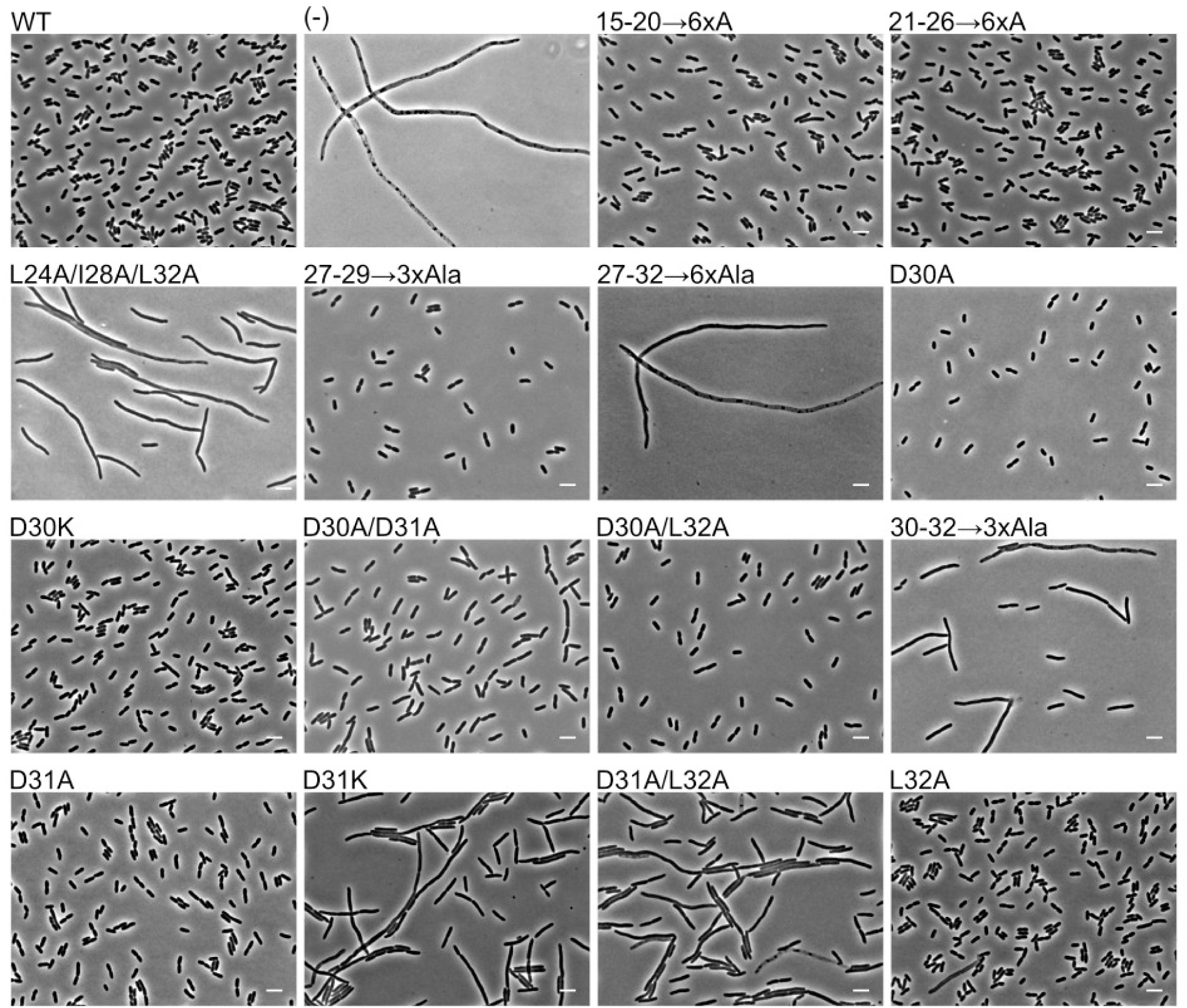
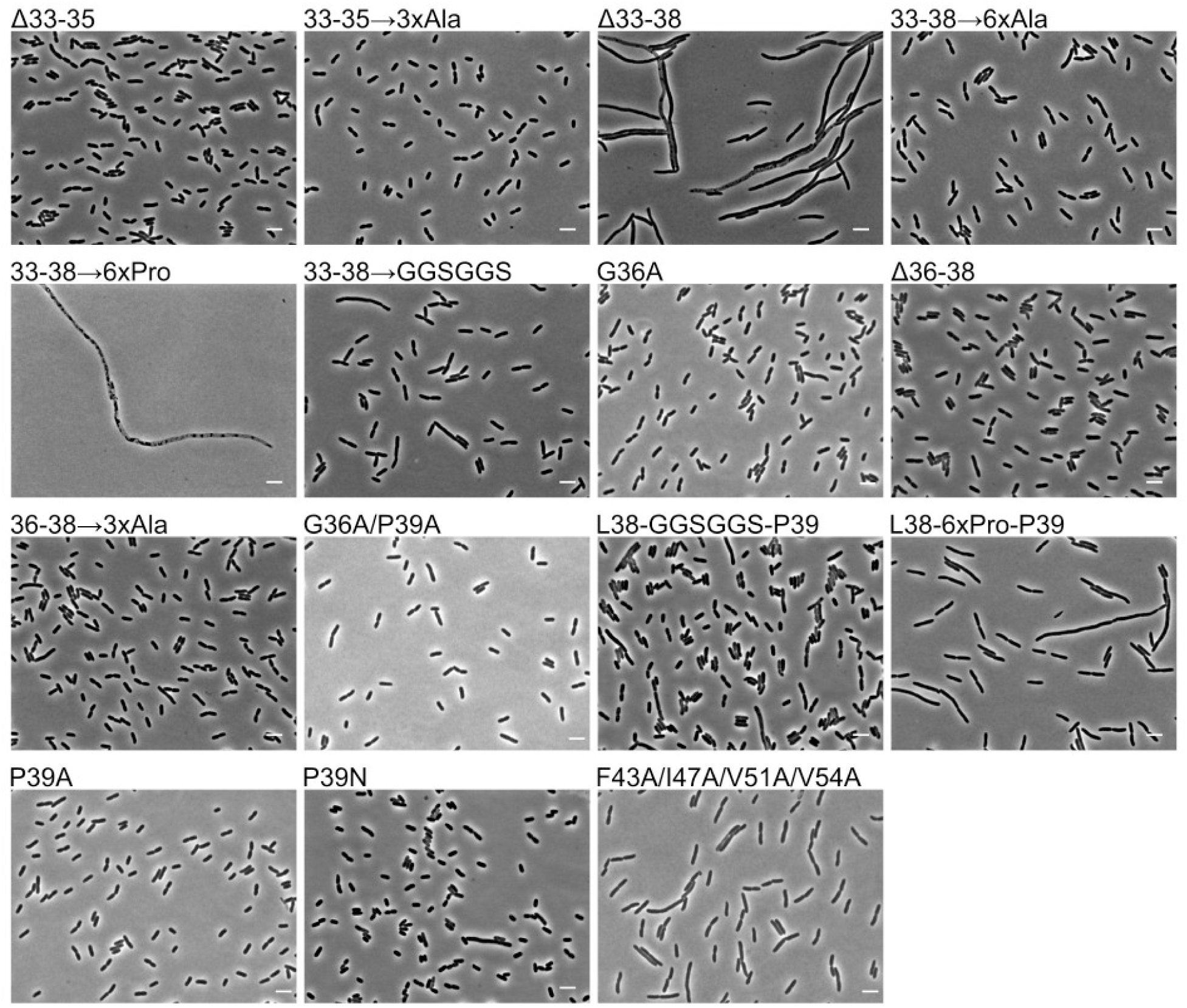
Representative phase-contrast images of *E. coli* cells expressing mutant variants of FtsL. Scale bar: 5 μM. Continues on the next page.

**Fig. S3.**
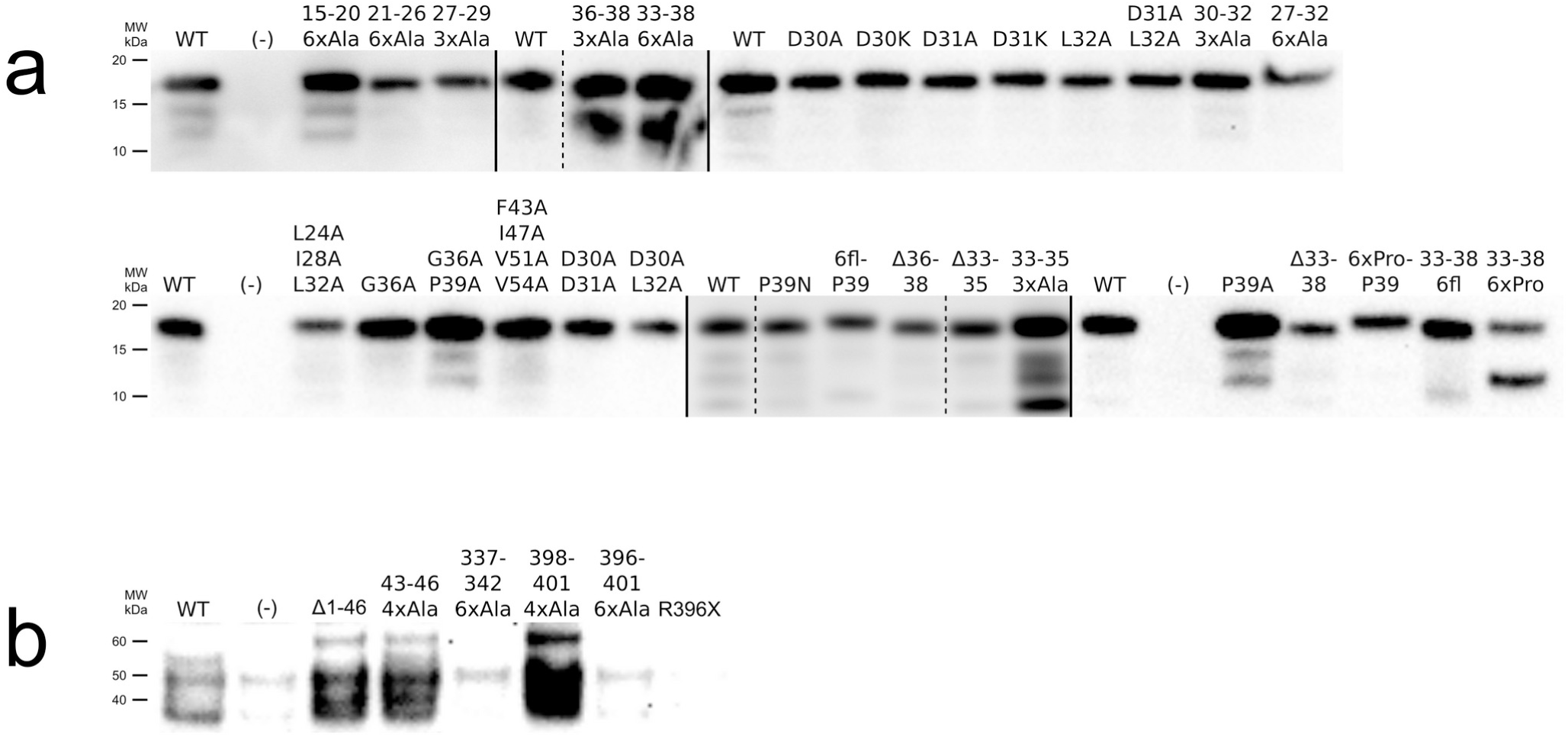
Expression level of FtsL and FtsW mutants assessed by western blot analysis. (a) Western blots for all of the FtsL mutants (anti-FLAG) tested in this work. (b) Western blots for select FtsW mutants (anti-GFP) from this work. FtsW samples could not be boiled (to enable anti-GFP binding), which likely contributes to the presence of multiple bands, though breakdown products are also a possibility. All samples were normalized to total protein concentration (determined by BCA assay). Protein expression level of the mutants with defective phenotypes are generally comparable to the respective wild type (WT). Negative controls (−) are from cells containing an empty vector expressing no FLAG- or GFP-tagged proteins. “6fl” refers to a 6-residue flexible sequence of GGSGGS. There are cases of mutants with increased breakdown product, though the abundance of the full protein is generally still comparable to WT. Splice sites within a single gel are separated by dashed lines (removal of duplicate samples), whereas individual gels are separated by solid lines.

**Fig. S4.**
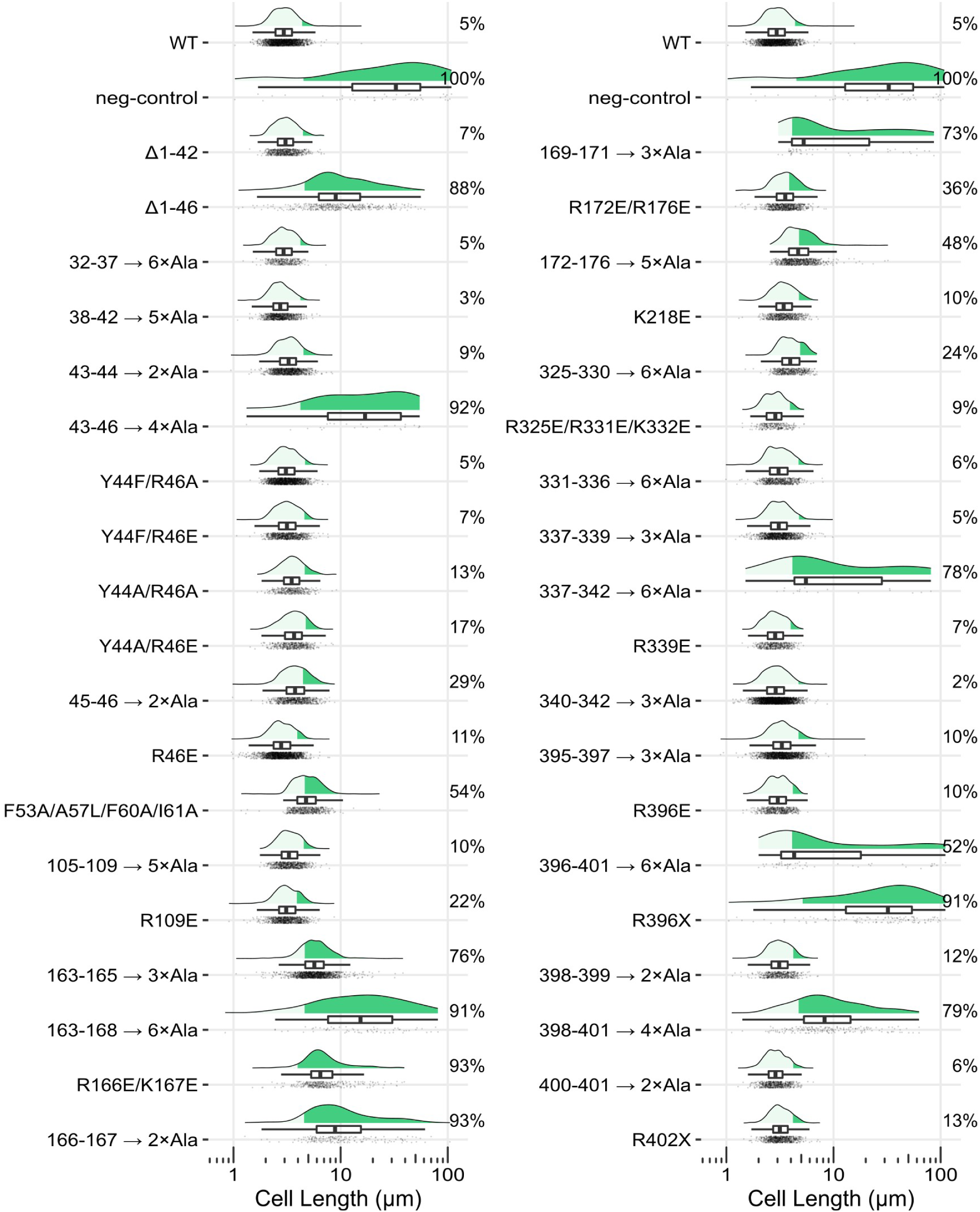
Effect of FtsW cytoplasmic tail and TM1 mutations. Summary of elongation defects for FtsW mutations tested within this paper. Elongated cells are defined as those exceeding the length of the 95^th^ percentile in the WT length distribution (green area; percentage indicates fraction of total population that is elongated).

**Fig. S5.**
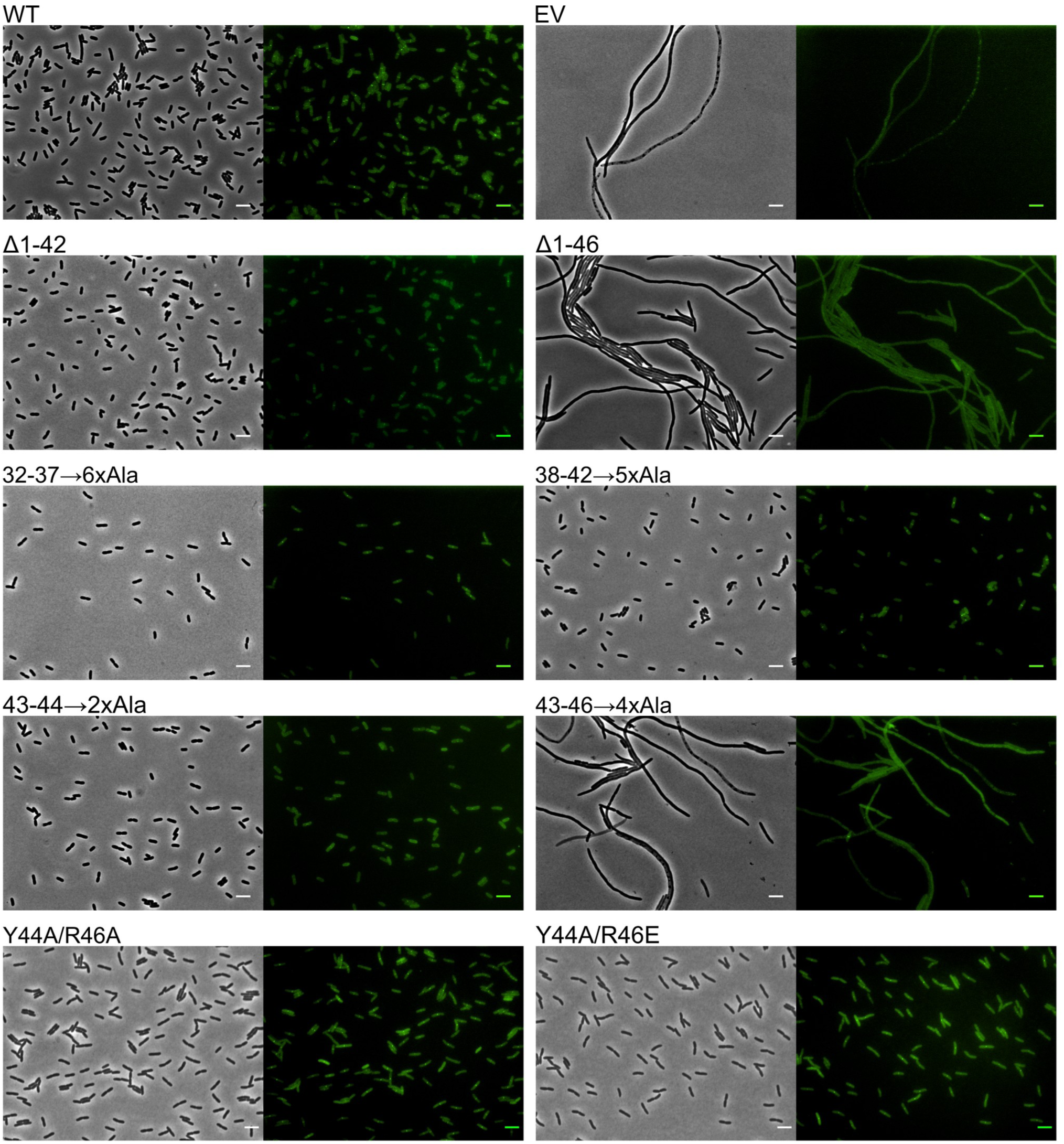
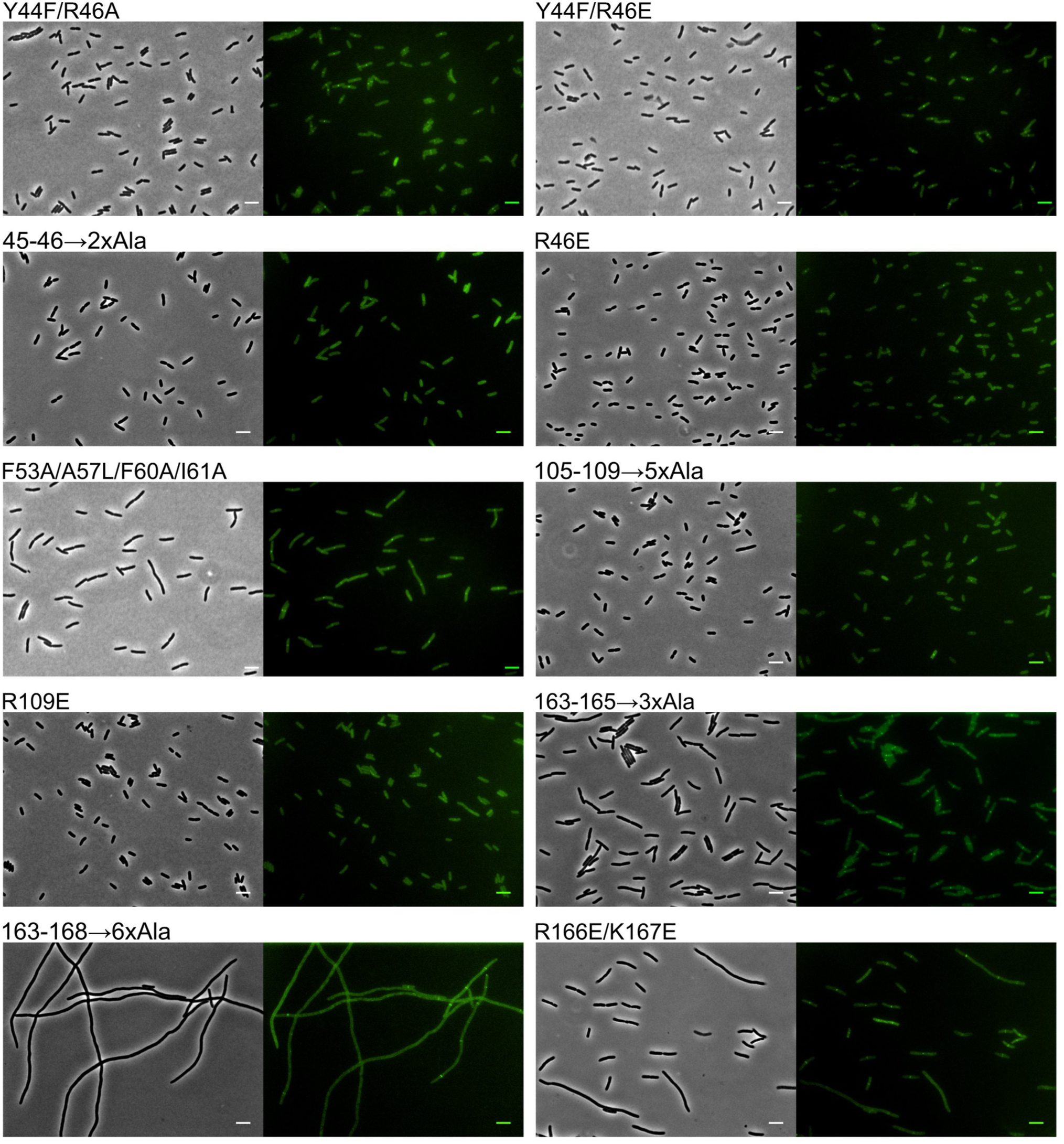
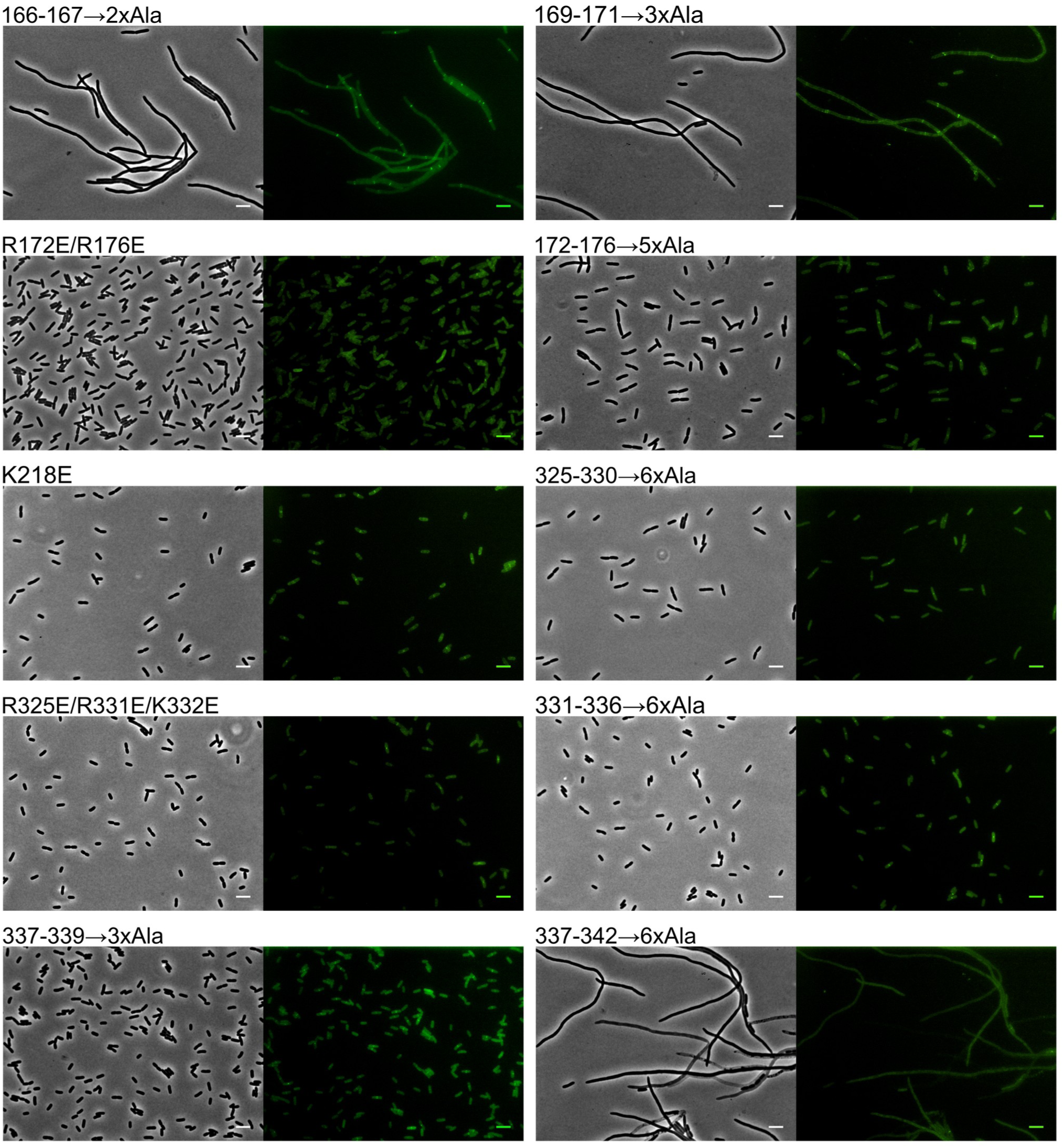
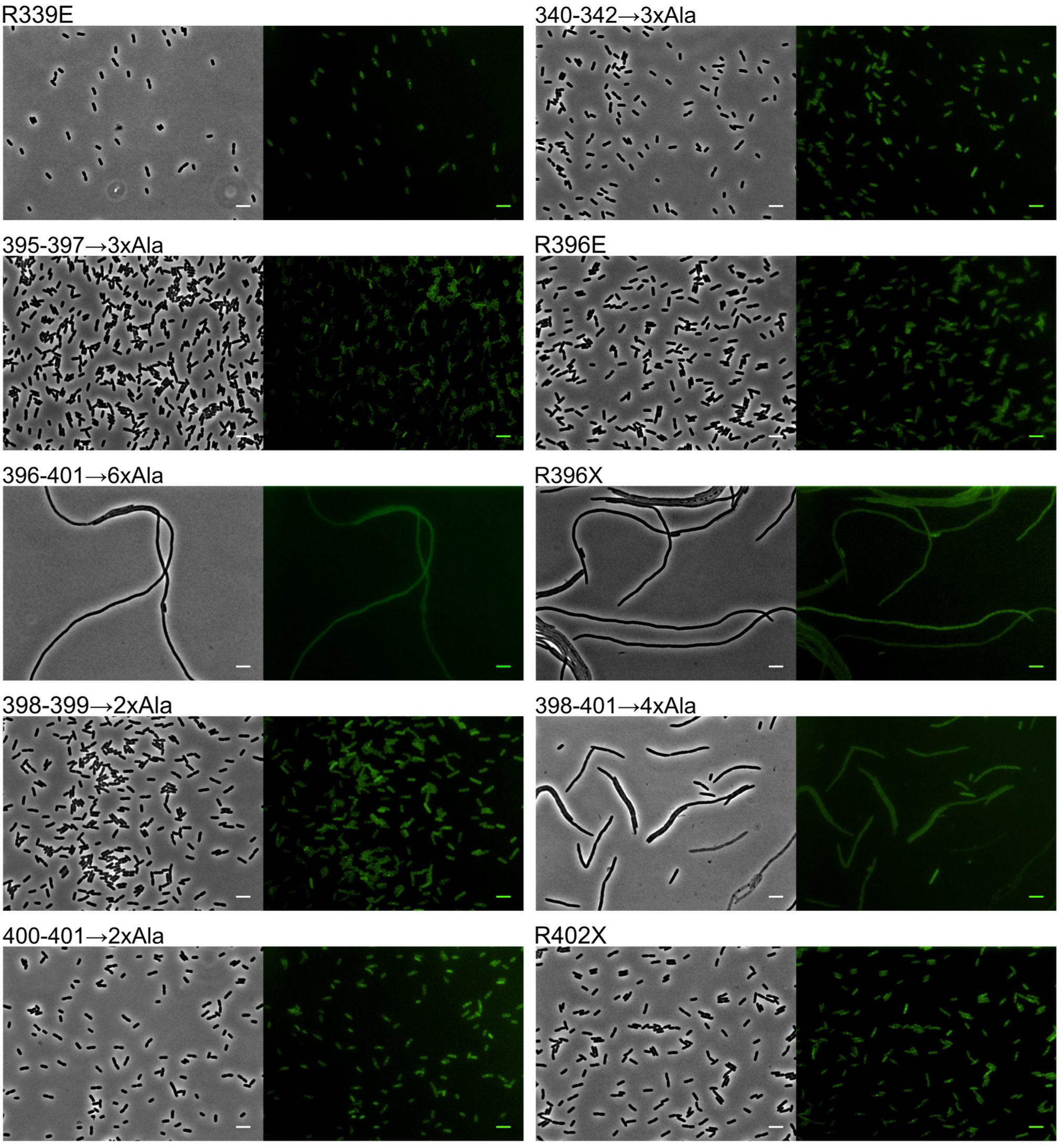
Representative phase-contrast (left) and GFP fluorescence (right) images of *E. coli* cells expressing mutant variants of FtsW. Scale bar: 5 μM. Continues on the next page.

**Fig. S6.**
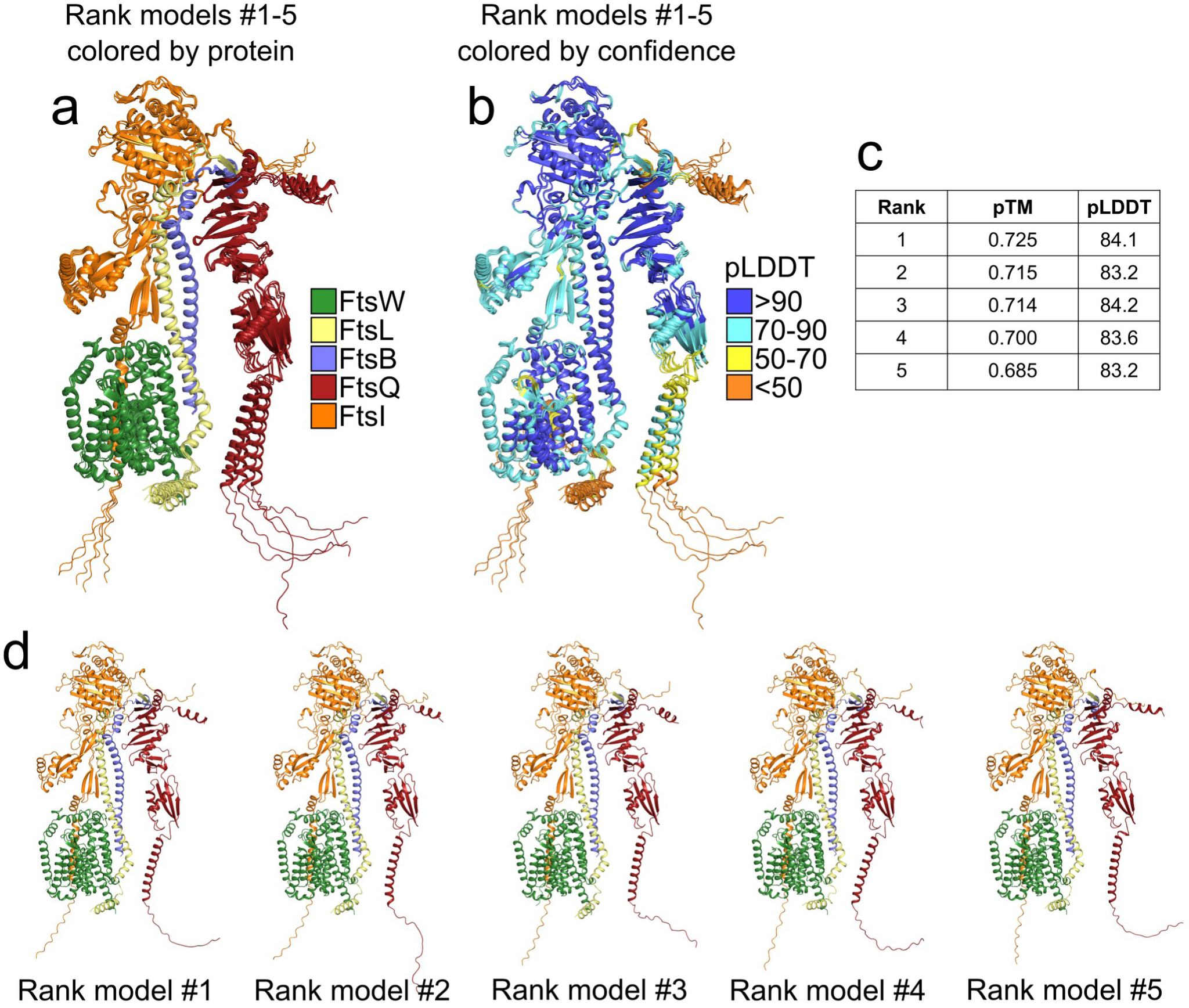
AlphaFold2 prediction of the FtsQLBWI complex. a) View of the five predicted structures superimposed and colored by chain. b) Same view, colored by the per-residue predicted LDDT score (pLDDT)^1^, which is a measure of confidence in the local structure surrounding the residue (blue: most confident; orange: least confident). Apart from the termini of the chains which have very low pLDDT scores, the five predicted structures align very closely with each other. c) The overall confidence of each predicted structure was estimated using a per-residue confidence metric called predicted local distance difference test (pLDDT) on a scale from 0 (least confident, orange) to 100 (most confident, blue) and the predicted TM score (pTM), which is a measure of similarity between two protein structures^2^. Structures were ranked by the pTM score. d) Views of the five individual models. Models are available for download at the Dryad Digital Repository (DOI TBD).

**Fig. S7.**
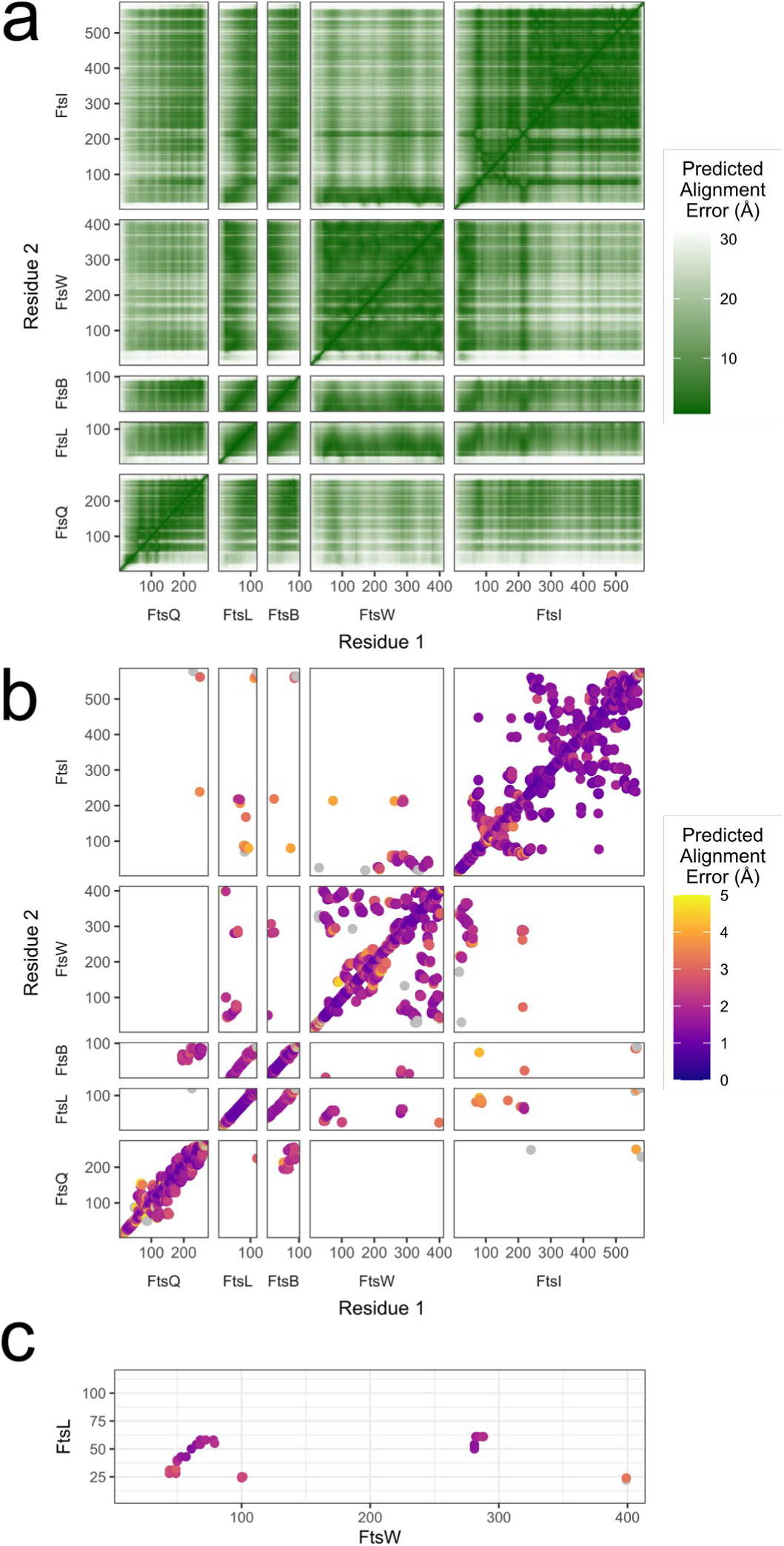
a) Predicted Alignment Error (PAE) plot of the top-ranked FtsQLBWI models predicted by AlphaFold2. b) PAE plot of contacting residues (minimum heavy atom distance < 4 Å) within the FtsQLBWI complex. Pairs of interacting residues with a PAE greater than 5 Å are colored gray. c) PAE of FtsL residues when using FtsW residues for the alignment. Color scheme is the same as b).

**Fig. S8.**
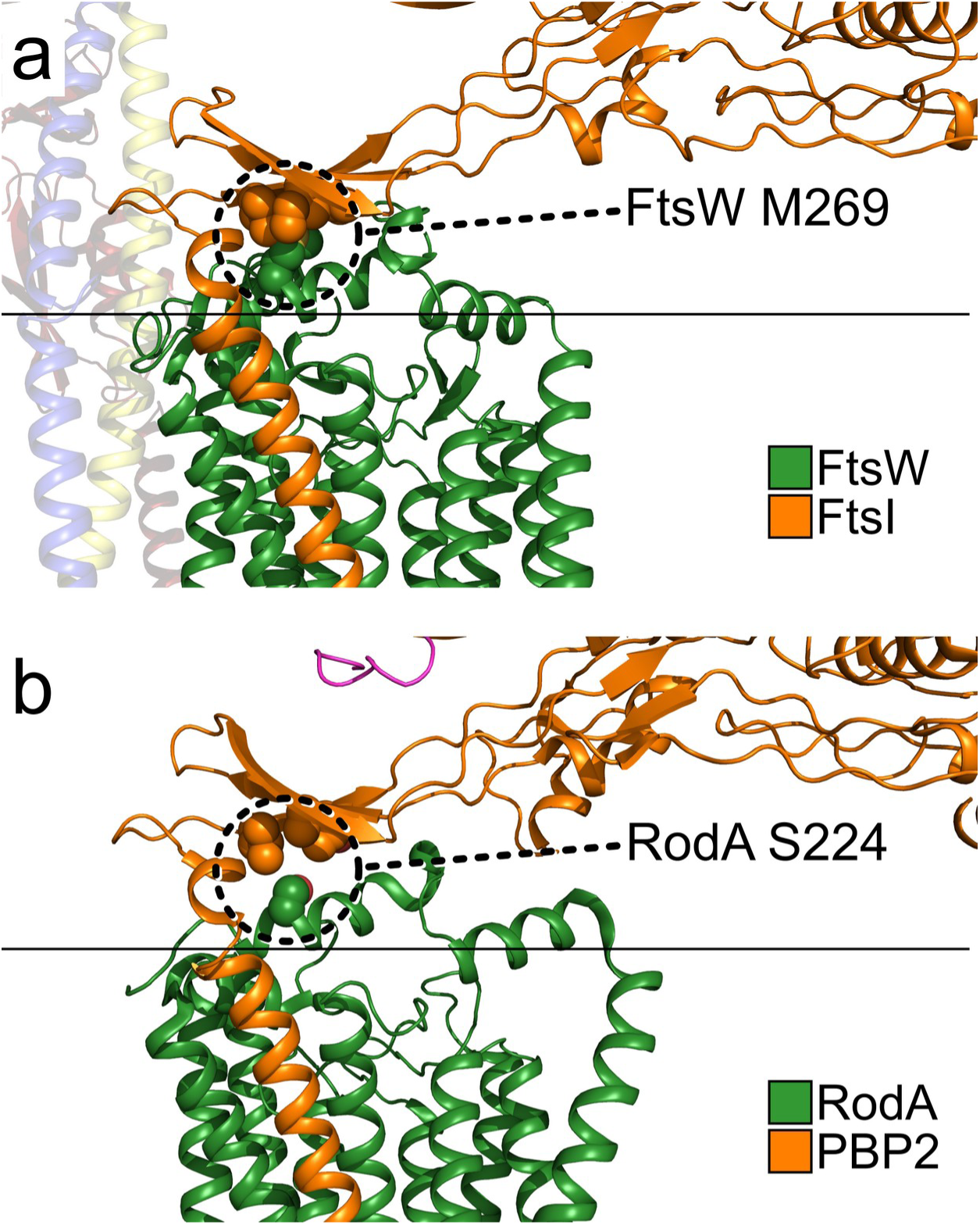
Interaction between the FtsW 7/8 loop and the FtsI pedestal domain. a) The interaction in the homology-based model appears to be mediated by FtsW M269, which forms a tightly packed, hydrophobic core with three FtsI residues (L62, V64, and I208). This position corresponds to the known FtsW superfission mutant M269I. b) Notably, this tightly packed, hydrophobic interaction is not observed in the original structural template of RodA-PBP2, in which M269 corresponds to a smaller and less hydrophobic serine residue S224.

**Fig. S9.**
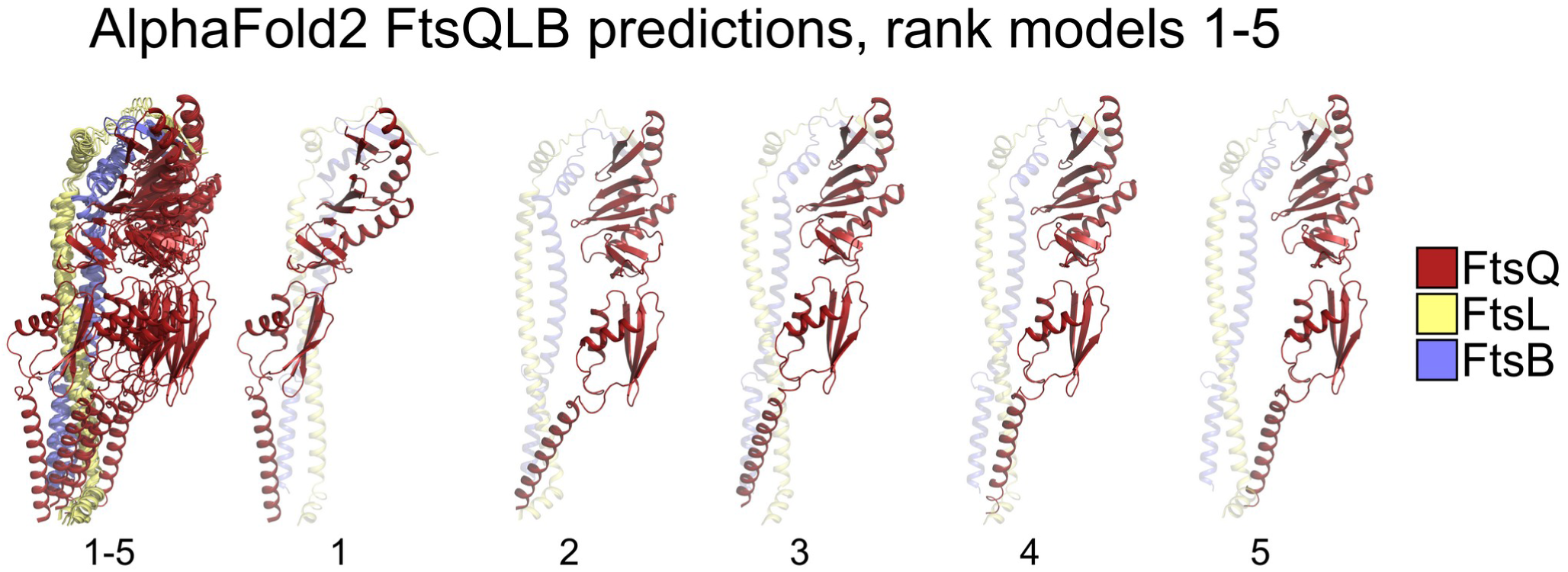
Configuration of FtsQ with respect to FtsLB in the AlphaFold2 predictions of FtsQLB. In the AlphaFold2 prediction of FtsQLBWI (supplementary Fig. S6), the FtsQ subunit assumes the same spatial orientation in all five top models. However, when the FtsQLB ternary complex is predicted by AlphaFold2, the resulting models display a wide variability in the spatial orientation of FtsQ with respect to FtsLB.

